# Topological defects drive influenza glycoprotein lattice assembly on spherical membranes

**DOI:** 10.1101/2025.11.06.686978

**Authors:** Zhehui B. Liu, Steinar Halldorsson, Thomas Calcraft, Lesley J. Calder, Peter B. Rosenthal

## Abstract

Lipid-enveloped viruses, such as influenza virus, assemble by budding from infected cell membranes, packaging internal components including the genome and acquiring an envelope containing surface glycoproteins in the process. Influenza C virus possesses a single surface glycoprotein, the haemagglutinin-esterase-fusion (HEF) protein that forms hexagonal arrays^1,2^ on the membrane envelope and is sufficient for budding of spherical particles^3^. However, a two-dimensional hexagonal lattice cannot completely cover a spherical virus membrane without defects. Using electron cryotomography (cryo-ET), we study the structural arrangement of the influenza C virus surface and find the hexagonal HEF lattice contains 5-fold and 7-fold defects organised in grain boundaries. The number of excess dislocations increases with system size while maintaining a net topological charge near 12. Our observations of defects in spherical crystals on influenza C virus particles of varying radius and shape matches theoretical predictions of continuum elastic theory^4^ for the proliferation of defects on soft lattices and experimental observations^5^ on colloidal systems. These findings provide new principles for assembly of pleomorphic viruses, extending the description of defects required for viral lattice assembly beyond the Caspar-Klug theory^6^ developed for isometric viruses. Our study informs a wide range of molecular self-assembly processes in biology and may also have implications for developing lattice materials with curved surfaces.

## Introduction

The haemagglutinin-esterase-fusion (HEF) envelope glycoprotein is a homotrimer that performs several essential functions in the influenza C virus life-cycle: it binds cell surface receptors at the start of infection, mediates fusion of the virus membrane with host membranes at the low pH of the endosomal compartment after internalisation into cells, and has a receptor-destroying enzyme activity that facilitates release of the virus from cells^7,8^. In addition, HEF trimers form a hexagonal 2D lattice on the virus surface^1,2^. The virus has a dense matrix layer beneath the membrane formed by the M1 protein and packages ribonucleoprotein particles containing the genome in its interior. M1 is a genetic determinant of virus morphology, and some mutations of the protein promote a filamentous particle shape for influenza C virus^9,10^. Virus infection also produces particles with hexagonal surface lattices that lack the matrix layer and internal components^3^, as is also the case with cellular expression of HEF alone^11^, indicating that the HEF glycoprotein in the membrane is sufficient to form the hexagonal surface lattice which mediates budding of spherical particles from cells.

However, 2D crystalline order as described for flat crystals cannot extend over the curved surface of a sphere due to geometric frustration. HEF lattice assembly therefore requires a more complete description consistent with spherical crystallography as well as the observed pleomorphy of influenza virus particles.

By contrast to influenza C virus, isometric viruses form spherical protein shells or capsids with icosahedral symmetry, typically with a specific number of copies of a single protein arranged as pentameric or hexameric capsomers which are the basic morphological units of the surface lattice. The organisation of capsomers into an icosahedral capsid in isometric viruses is described by Caspar-Klug theory^6^ and is based on triangulation of a hexagonal lattice, with a variable number of hexamers and the insertion of exactly 12 pentamers as defects that introduce curvature to the flat hexagonal lattice to form a closed shell (**Fig. 1a**). The total number (*N*) of capsomers depends on the triangulation number *T* (*N* = 10*T* + 2, where *T* = *h*^2^ + *hk* + *k*^2^, and *h* and *k* are integers ≥ 0). A further generalisation describes HIV capsids^12–14^ that are organised as a hexagonal lattice but with 12 pentamers arranged to enclose a shell with fullerene cone geometry. Icosahedral capsids are also used to package nucleic acid within lipid enveloped viruses^15^ and in other cases form shells to enclose and shape viral membranes^16^.

**Fig. 1.**
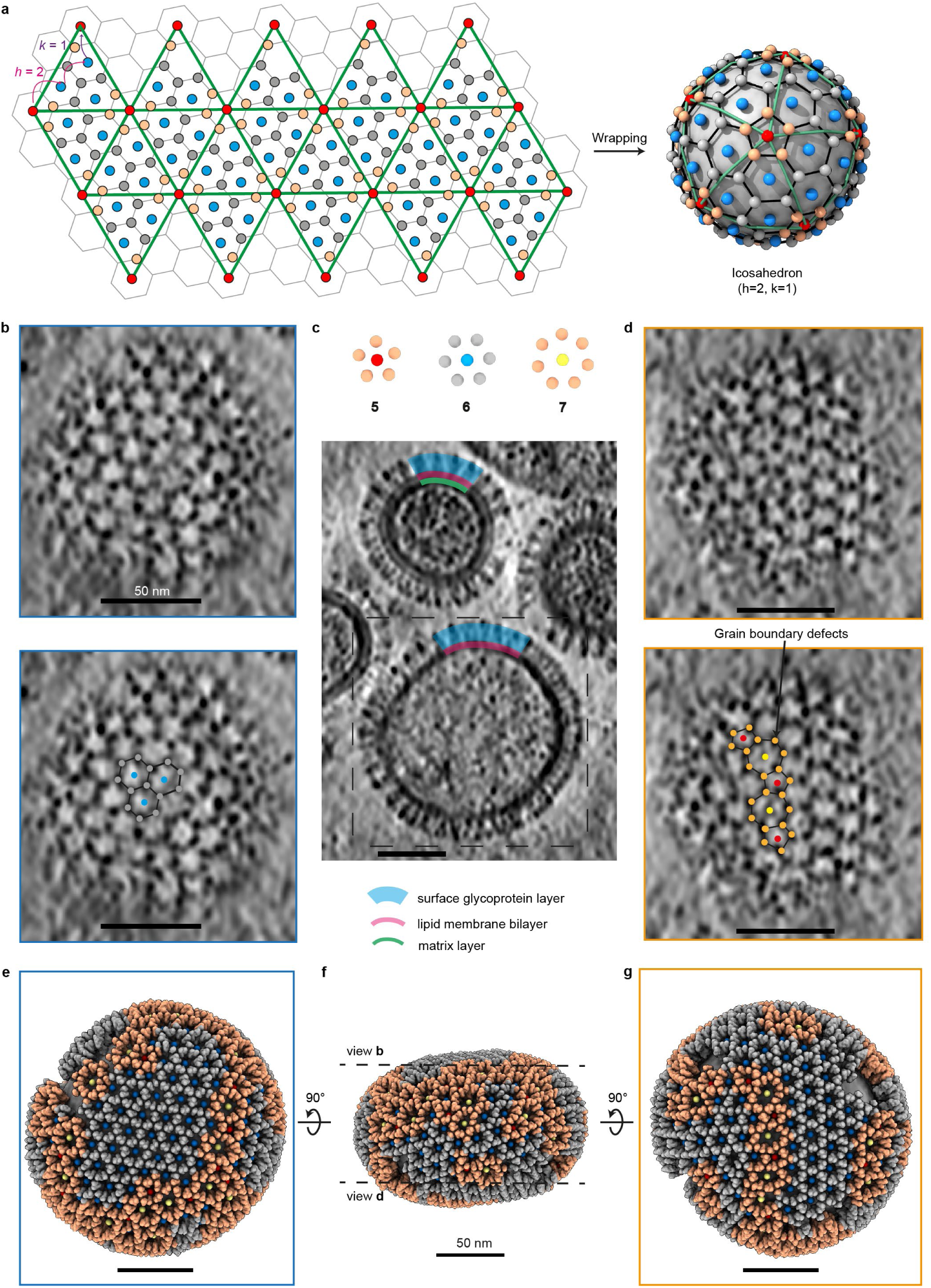
Cryo-ET reconstructed tomogram and modelled HEF surface lattice on membrane. **a,** Schematic showing that a flat 2D hexagonal lattice (left) requires defects to wrap a sphere (right). As described by Caspar-Klug theory, triangulation (green) of the 2D lattice (formed of grey and orange spheres as lattice vertices and blues spheres as hexagon centres) and insertion of 12 pentagonal defects or disclinations (at red marker locations) provides a covering of the sphere with icosahedral symmetry. In this example the triangulation number *T* = 7 (*h* = 2, *k* = 1). **b-d**, 3D tomogram of a single virus particle (**Extended Data Table 2**, virus particle no. 24). **b**) top, tomogram section through surface shows HEF glycoprotein trimers (black densities) forming a lattice on the virus surface; bottom, identical section as above with hexagonal arrangement of several HEF trimers indicated (each grey marker sphere denotes an HEF trimer and each blue marker sphere represents the centre of a hexameric arrangement). **c**) tomogram section with a larger field of view through the middle of the virus particle shows that it lacks a matrix layer in contrast to an adjacent virus particle with a matrix layer and denser content in interior (example segments of surface glycoprotein layer, lipid membrane bilayer and matrix layer are coloured in blue, pink and green). **d**) top, tomogram section through the surface on the opposite side of the virus particle; bottom, identical section where several HEF trimers (orange marker spheres) are labelled where they form 5-fold and 7-fold defects arranged in a grain boundary; 5-fold defects are pentagons with orange HEF trimers at vertices and a red marker at centre and 7-fold defects are heptagons with orange HEF trimers at vertices and a yellow marker at centre. **e-g,** Spherical lattice model of the virus particle with HEF trimers distinguishing areas by grain boundaries (orange) and regular hexagons (blue). **e)** same view as **b**, showing a central area of hexagons on the flatter surface. **f)** a side view displaying defects clustered in highly curved areas, with detailed curvature analysis using the same virus particle as an example in **Extended Data** Fig. 8. Location of tomogram sections in **b** and **d**, indicated. **g)** same view as **d**, featuring “5-7” defect chains. Note: The blue, red and yellow round markers are symbolic, indicating the centres of hexagons, 5-fold defects, and 7-fold defects, respectively, correlating with direct observations from the tomogram. Scale bar: 50 nm.

Beyond viruses, lattice defects determine the physical properties of crystalline materials, carbon fullerenes^17,18^, nanotubes^19^ and soft matter materials. Defects observed in hexagonal lattices may be characterised by assigning a topological charge (*q* = 6 - *n*), which is defined by the deviation of its local coordination from 6, where *n* is the local coordination number. An isolated 5-fold or pentameric (7-fold or heptameric) defect, known as a disclination, introduces a local angular deficit (excess) of 2π/6 and has a charge of +1 (−1), whereas an interacting “5-7” defect pair, known as a dislocation, has a net charge of 0 as do the neutral hexagons that form the bulk of the lattice. The topology of a sphere or closed surface (Euler characteristic χ = 2) requires a total angular deficit of 4π (Gauss-Bonnet theorem)^20^, corresponding to a topological charge^21^ of 12, as observed for the icosahedron.

The number and arrangement of defects which minimise the energy of a spherical crystal lattice is described by continuum elastic theory^4,22^. This approach considers the mutual interaction of the defects alone within the surrounding elastic medium without requiring a microscopic description of the interactions and reduces the degrees of freedom in the analysis of this complex problem. Finding the optimal arrangement of interacting defects on a sphere is related to the Thomson problem^22–24^ of identifying the minimum energy configuration of electrons in atomic shells. Continuum elastic theory has been applied to explain defects in various systems, including colloidosomes^25,26^, icosahedral virus capsids^27,28^, and elastomers^29^. Compared to isometric virus capsids, influenza C virus is pleomorphic and envelope proteins on the membrane surface form a weakly-associated 2D lattice that displays behaviours similar to those of other soft matter systems.

To understand how HEF drives virus assembly and how it can form a surface lattice, we visualise and model the surface lattice by electron cryotomography (cryo-ET)^30^ to characterise the arrangement of the glycoproteins on the surface of influenza C virus particles. The average system size of the spherical lattice (*R/a*), defined as the ratio of the virus radius *R* to the lattice spacing *a*, is typically greater than 5 (∼ 450 Å / 85 Å), which is the critical size at which strings of dislocations called grain boundaries form^4^. We observe grain boundaries and characterise the defect arrangements on virus particles with close-to-full spherical coverage by the crystalline lattice, revealing an alternative system for defect-mediated pleomorphic virus assembly to the icosahedral geometry described by Caspar-Klug theory. The findings will extend current soft-lattice design approaches to molecular packaging and cargo delivery.

## Results and Discussion

### HEF envelope glycoprotein lattice exhibits grain boundaries

We employed cryo-ET to reconstruct spheroidal influenza C virus particles in three dimensions. The 3D reconstructions revealed HEF spike glycoproteins in the membrane envelope arranged in an open hexagonal lattice mediated by dimeric interactions between trimers as previously described^1,3^. Notably, the analysis revealed 5-fold and 7-fold defects organised as grain boundaries, features directly observable in the tomograms (**Fig. 1b-d**). Such defects were observed in all virus particles in three datasets for a variety of sizes and morphology (**Extended Data Table 1,** >1000 individual virus particles).

Subtomogram averaging (STA) obtained reconstructions of individual trimers and three types of oligomeric assemblies: a hexamer of trimers, a pentamer of trimers, and a heptamer of trimers (**Fig. 2a-d** and **Extended Data** Fig. 1). Subtomogram averaging of particles from the pentamer and the heptamer, reconstructed at a larger size, show a composite oligomer, consisting of 5-fold and 7-fold defects bound together as “5-7” pair (**Fig. 2e**). This “5-7” dislocation map shows densities of all three types of oligomeric assemblies associated together, as a key component of the surface lattice. These oligomeric forms are the basic morphological units of the HEF lattice analogous to capsomers, with the pentamers and heptamers of trimers forming 5-fold and 7-fold defects within the lattice formed of hexamers of trimers.

**Fig. 2.**
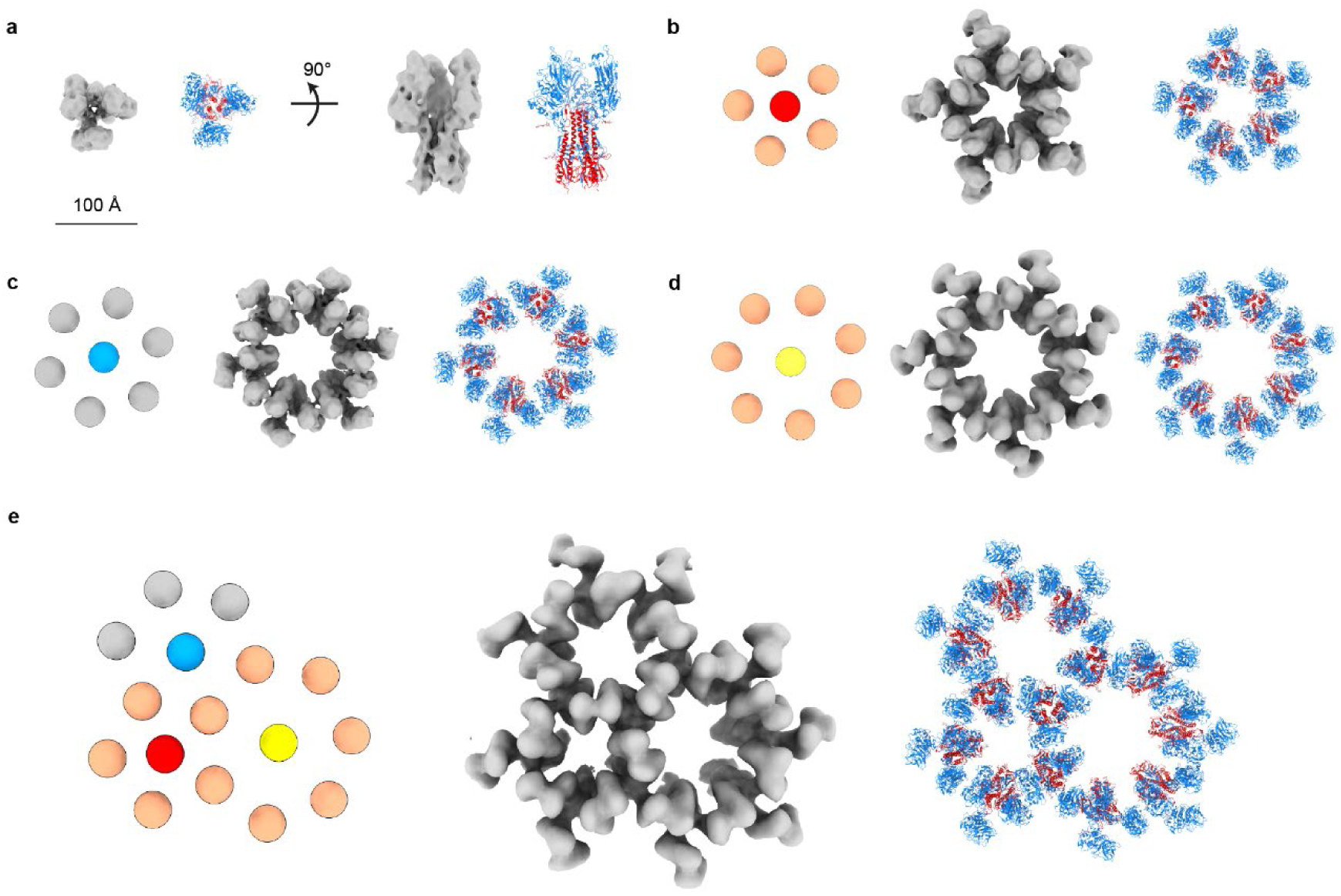
Subtomogram average (STA) maps of HEF glycoprotein in different oligomeric configurations. STA maps of HEF trimers (grey map) fitted with HEF models (PDB: 6YI5; with HEF1 subunit in blue and HEF2 subunit in red). **a,** Single HEF trimer, resolution: 7.4 Å, in both top view and side view. Scale bar: 100 Å. **b,** Pentameric assembly of HEF trimers shown schematically (orange spheres for HEF trimers; red sphere indicating the pentamer centre), as STA map (12.2 Å) and as fitted model. **c,** Hexameric assembly of HEF trimers shown schematically (blue spheres), as STA map (9.1 Å) and as fitted model. **d,** Heptameric assembly of HEF trimers shown schematically (orange spheres for HEF trimers; yellow sphere indicating the heptamer centre), as STA map (19.6 Å) and as fitted model. **e,** Composite oligomer comprising a dislocation formed by pentamer and heptamer defect oligomers and adjacent hexamer, shown schematically (orange spheres for the defect assemblies; blue spheres for the hexamer unit; red and yellow spheres marking the centre of 5-fold and 7-fold defects), as STA map (27.2 Å) and as fitted model.

Our previously described in situ HEF trimer structure^3^ (PDB:6YI5), different from the X-ray crystal structure^7^, has a splayed conformation that mediates lattice contacts. We fit this structure to STA reconstructions to construct models for the three types of oligomer (**Fig. 2b-d**) and the composite oligomer containing the “5-7” dislocation pair (**Fig. 2e**). The interaction between HEF trimers in the lattice occurs at the head-to-head dimeric interface formed by two surface loops^3^ in the hexamer (**Extended Data** Fig. 2).

We modelled the glycoprotein layer on the virus particle surface by an approach based on template matching, using all three types of oligomer STA maps as search objects, initially assigning oligomer type and centre positions on the surface (see **Methods, Extended Data** Fig. 3). Positional and orientational information for individual glycoprotein trimers were assigned to the highest trimer correlation peak near an assigned hexamer vertex, which also assigns shared trimers to the adjacent defect oligomers (**Extended Data** Fig. 3). Subsequently, the identification of oligomer types was verified or reassigned by counting the coordination number of glycoprotein vertices enclosing each polygon (**Extended Data** Fig. 4). The assignment of modelled oligomer type and orientation was also verified by performing subtomogram averaging and back-plotting onto a membrane surface yielding a geometrically consistent lattice model in which positions of shared glycoprotein molecules between adjacent oligomers superimpose (**Extended Data** Fig. 5 and **Extended Data** Fig. 6).

Geometric analysis (see **Methods**) shows that the assigned oligomeric polygons are largely regular in shape, with hexamers being more regular than pentameric and heptameric defects as quantified by the normalised root-mean-square deviation (RMSD) of individual HEF trimers from the expected polygon radius: 72.3 Å for pentagons (RMSD: 6.8 Å), 85.0 Å for hexagons (RMSD: 6.5 Å), and 97.9 Å for heptagons (RMSD: 10.1 Å).

An example of a virus particle surface model and its corresponding tomogram features is shown in **Supplementary Movie 1**. Throughout the analysed influenza C virus particles, which range in radius from approximately 35 nm to 80 nm (**Fig. 3**), the observed arrangements include 5-fold disclinations and grain boundary defects that mainly form in linear chains of dislocations in “5-7-5” arrangements surrounded by hexagonal lattice, but also as branched or joined grain boundaries (**Fig. 3** and **Extended Data** Fig. 7). The surface glycoprotein arrangement with topological defects is asymmetric and exhibits no global point symmetry.

**Fig. 3.**
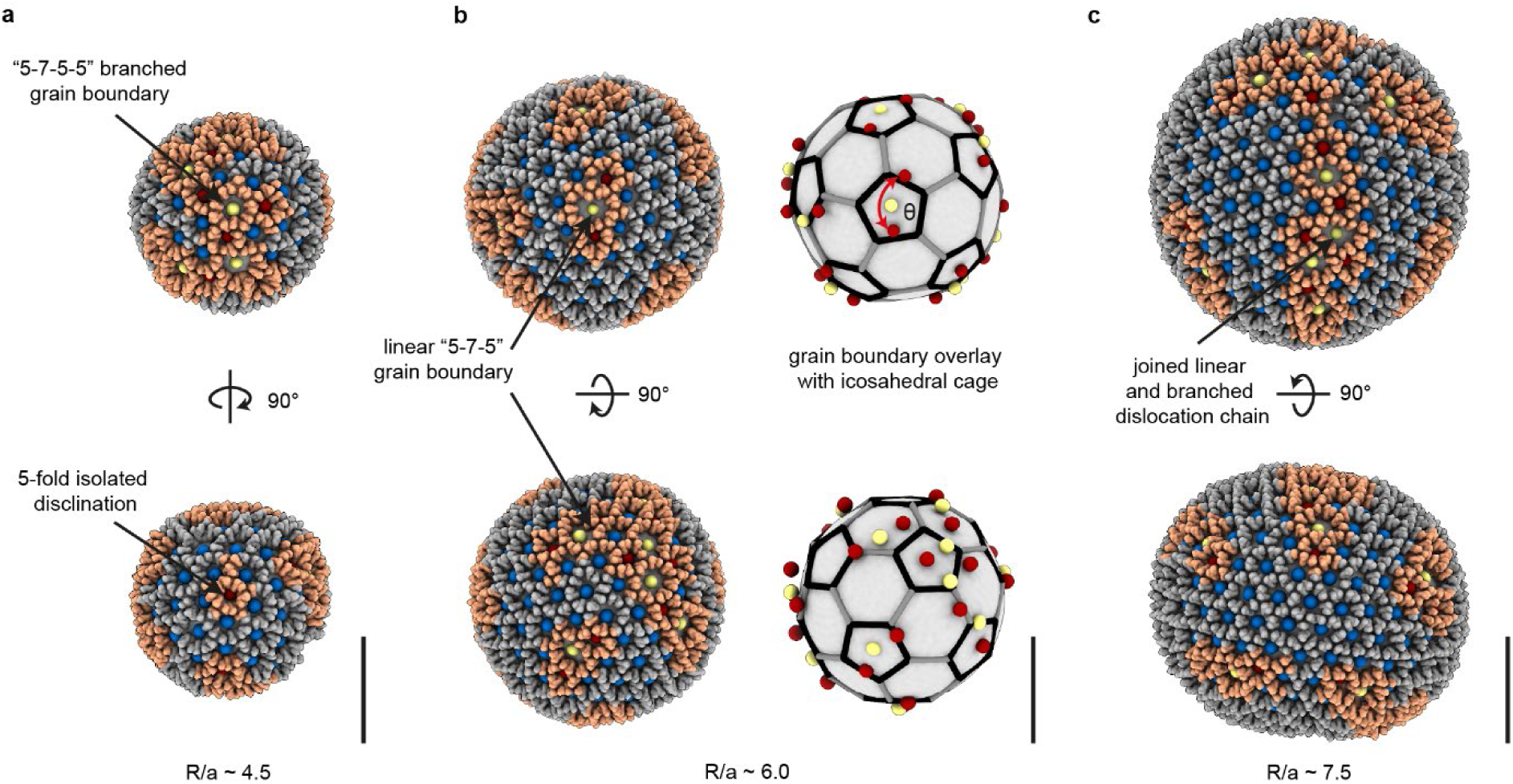
Topological defects in HEF lattices on spheroidal virus particles of varying size. Representative examples of three virus particles of different sizes. **a,** A small virus particle (**Extended Data Table 2**, no. 16) with system size *R/a* ≈ 4.5, where *R* is the area-equivalent radius 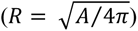 and *a* ≈ 8.5 nm is the spacing between adjacent HEF trimers. Both isolated 5-fold disclination and branched “5-7-5-5” dislocation defects are seen. **b,** A medium-sized virus particle (no. 29) with *R/a* ≈ 6.0, with linear “5-7-5” grain boundaries whose arrangement is compared to an icosahedral cage, with good matches in certain orientations but not all. **c,** A large virus particle (no. 42) with *R/a* ≈ 7.5. This virus particle is slightly ellipsoidal (aspect ratio ≈ 1.2). Linear and branched dislocation chains are joined to form longer grain boundaries. Colour coding follows that used in Fig.1. Scale bar: 50 nm.

### Spherical crystallography of influenza C virus

The HEF lattice on the membrane surface of influenza C virus has a significantly different organisation from icosahedral virus capsids. The differences correspond to two types of predicted behaviour for spherical crystals depending on the energetic cost of introducing defects. In continuum elastic theory, the energy of the spherical crystals has contributions from interactions between disclinations, dislocations, and their interactions with the curved medium, the latter expressed as 𝐺𝐺(𝑥𝑥) − ∑_𝛼𝛼_ 𝑞𝑞_𝛼𝛼_ 𝛿𝛿(𝑥𝑥 − 𝑥𝑥_𝛼𝛼_), where *G(x)* is the Gaussian curvature of the surface and *q_α_* are topological charges^4,26^. Topological charges are therefore arranged to minimise repulsive interactions between disclinations while being attracted to regions of same-sign curvature. The curvature can be thought of as screening repulsive interaction between isolated disclinations.

Application of the elastic theory to small virus capsids predicts that the distribution of 12 isolated 5-fold disclinations at the vertices of an icosahedron minimises the energy^27,31–33^ while satisfying the topological charge requirement. The introduction of an isolated disclination has a considerable energetic cost particularly at large system sizes where the disclinations are not adequately screened due to lower Gaussian curvature. The energy of the disclination can be relieved by buckling of the lattice, with the threshold^21,27,34^ for buckling given by the Föppl– von Kármán (FvK) number, 𝛾𝛾 = 𝑌𝑌𝑅𝑅^2^/𝜅𝜅, where *Y* is the Young’s modulus and *κ* is the bending rigidity. Thus, for an icosahedral capsid with a large Young’s modulus and large system size such that FvK > 154, the lattice buckles to form faceted icosahedral structures as observed for large capsids, unless the capsid shell is assembled around a scaffolding protein^35^.

In contrast to virus capsids, the HEF trimers constrained to the membrane surface make weak interactions based on the small contact points and small interface binding energies (∼2 kcal/mol, **Extended Data** Fig. 2) and virus particles remain approximately spheroidal without regular faceting of the lattice and membrane. For low Young’s modulus and low FvK number, the energetic cost of additional neutral charge defects is lower and may be preferred over buckling^4,21^.

When the system size (*R/a*) becomes sufficiently large, its Gaussian curvature decreases and is insufficient to screen the long-range repulsive interactions of isolated disclinations. Continuum elastic theory predicts that additional dislocations carrying neutral topological charge (excess defects in “5-7” pairs or dipoles) are introduced to screen disclinations and reduce strain. Thus, the formation of grain boundaries^5^ is anticipated when the system size, *R/a,* reaches approximately 4-5, and the total number of particles 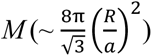, exceeds approximately 300 to 400^4,24,36^. We therefore characterise the influenza C virus surface lattice in the context of this general theory of defects.

### General properties of the observed grain boundary defects and the role of system size

We analysed influenza C virus particles with near-complete lattice coverage (mean coverage = 86 ± 7%) on their spherical membranes and built marker models for detailed defect analysis (*n* = 42 virus particles, **Extended Data Table 2**). When extrapolated to a complete lattice (observed charge / coverage fraction), most of these virus particles exhibit a topological charge near 12 (mean = 11.41) in accordance with Euler’s theorem, despite variation in size or morphology (**Fig. 4a** and **Extended Data Table 2**). The number of defects typically exceeds the minimal 12 pentamer requirement by several fold. Our tomographic analysis of influenza C virus particles is based on more complete lattice coverages than earlier experimental measurements^5,37^ of total topological charge in soft matter systems, such as colloidal particles in a water-in-oil emulsion system, where lattice coverages are of less than 50%. The slight deviations from a topological charge of 12 on these nearly complete lattices may be due in some cases to voids^38^ in the lattice, or loss of lattice areas during specimen preparation or from imperfect reconstructions due to noise and the missing wedge problem in tomography (anisotropic data due to limited tilt range) which can lead to inaccurate defect identification. While it seems unlikely that a perfect spherical crystalline surface lattice with a precise topological charge is a necessary condition for virus assembly, the observation demonstrate that topological defects drive the formation of continuous lattices covering virus particle surfaces.

**Fig. 4.**
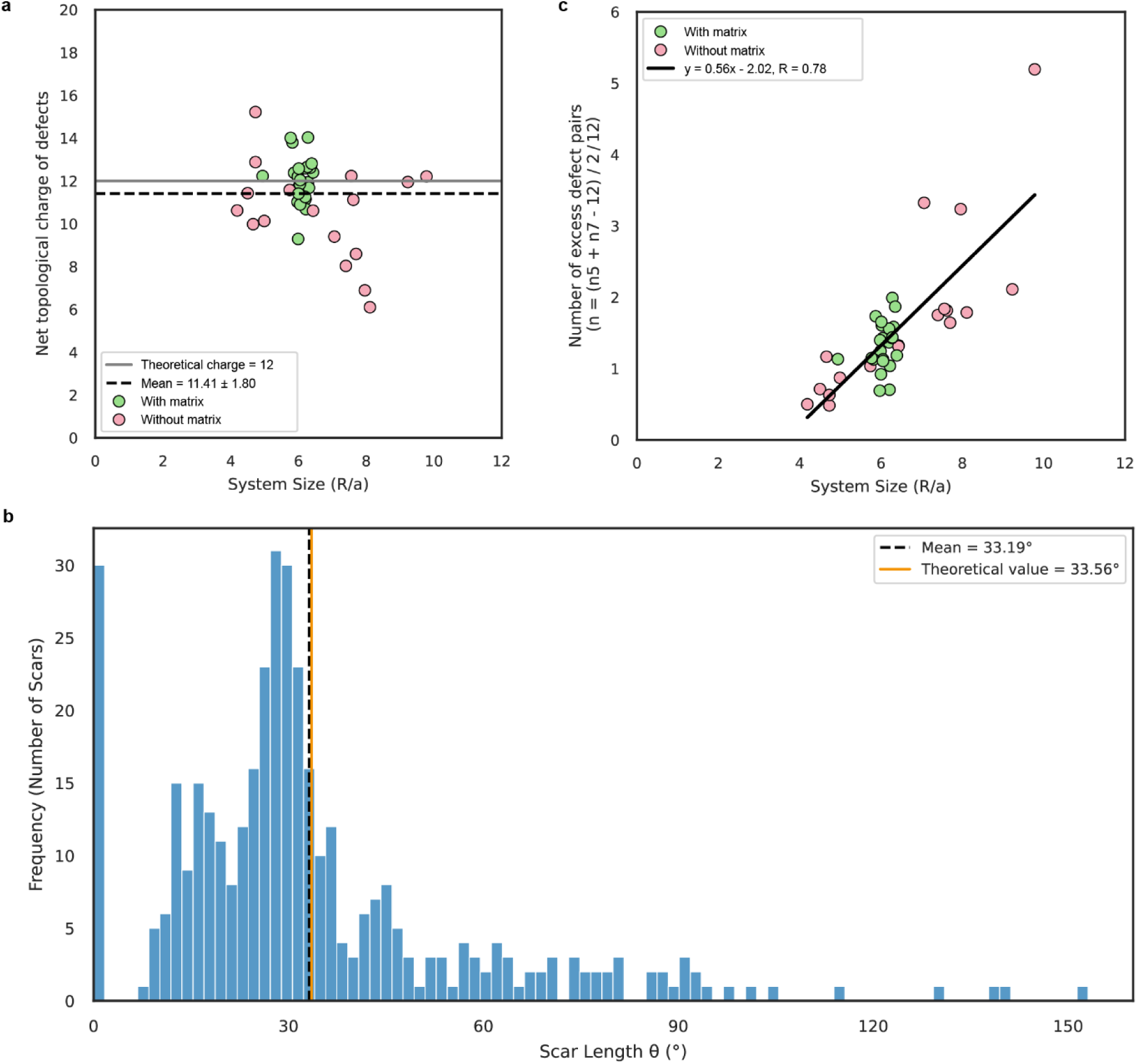
Topological charge and analysis of grain boundary defects with system size. **a,** The total topological charge (difference between the number of 5-fold defects and 7-fold defects) of the lattice system in most virus particles is near 12, consistent with Euler’s theorem. The mean topological charge of the dataset is 11.41 ± 1.80 (*n* = 42), with total topological charge being extrapolated as described in **Methods** and **Extended Data Table 2**. **b,** Histogram of the angular lengths (θ in **Fig.3b**) of all grain boundaries. The mean length for all boundaries is 33.19° (dashed line), consistent with the theoretical prediction of 33.56° (solid line in orange). The distribution is broad (s.d. = 23.90°), featuring a peak at 0° corresponding to isolated 5-fold disclinations prevalent in smaller virions, and a clustering around ∼30° (median = 28.76°). When excluding these isolated defects, the mean length of scars containing dislocations is 35.99°. **c,** The number of excess defect pairs per 12 disclinations vs the system size *R/a* (slope = 0.56; Pearson correlation coefficient *r* = 0.78, *p* < 0.05). Matrix-containing virus particles (pink) cluster near R/a ≈ 6, whereas matrix-free particles (green) span a broader size range.

The average number of grain boundaries after extrapolation for coverage is 12.33 ± 3.36 (**Extended Data Table 2**), with each carrying a mean net charge of ∼1, consistent with grain boundaries screening isolated disclinations. These appear to be distributed isotropically but without strictly obeying icosahedral or other point symmetries (**Fig. 3b**). The length of the grain boundaries, or “scars”, defined as the angular extent of a scar from a terminating 5-fold disclination, spans a wide range from 0° (for an isolated disclination) to 150° but with a mean of 33.19° (s.d. = 23.90°) (**Fig. 4b**). This value is in agreement with continuum elastic theory which predicts 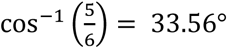, marking the termination point of dislocation growth^4,36^. Equivalently, the theory predicts a linear growth relationship between the number of excess defects and system size (*R/a*) at a rate of 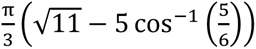, or 0.41. For influenza C virus particles, the number of excess defect pairs, *N* (per 12 disclinations) on spheroidal particles increases with *R/a* with a slope of 0.56 (**Fig. 4c** and **Extended Data Table 2**). Together, these observations reflect a universal behaviour for defects in a spherical crystal, independent of a microscopic description of molecular interactions, as also reported for colloidal systems^5^ on the micrometre scale and spherical elastomer crystals^29^ on the millimetre scale.

In modelled spheroidal particles, there is a clustering of more uniformly-sized particles at *R/a* ≈ 6 of particles with a matrix layer beneath the membrane (**Fig. 1c**, coloured in green) which may, along with packaged internal components, impose curvature of the particles. The small and large particles analysed typically lack a matrix layer (**Fig. 4a,c**). The observed small virus particles correspond to the theoretical threshold (4-5) at which grain boundary defects begin to emerge. Large virus particles (*R/a* ≈ 7-10) are generally ellipsoidal (**Extended Data Table 2**) and adopt a preferred orientation with their long axis in the plane of the ice film which may be the result of compression (**Fig.1** and **Extended Data** Fig. 8c).

The types of grain boundary motifs observed are more diverse than previously reported in experiments and simulations^4,5^. The linear “5-7-5” motif consisting of three defects (**Fig. 3b** and **Extended Data** Fig. 7c) is observed across all system sizes but are particularly prominent in mid-sized virus particles, often when the number of excess dislocation pairs is near one. In small virus particles, in addition to isolated 5-fold disclinations, a branched motif, typically with three 5-fold disclinations surrounding a central 7-fold disclination are observed (**Fig. 3a** and **Extended Data** Fig. 7f). This configuration potentially contributes a local topological charge of +2, satisfying the global topological constraint while keeping a low number of grain boundaries in small systems. The “5-7” dislocation pairs (**Extended Data** Fig. 7b), which carry zero net topological charge, are often adjacent to vacant areas where missing defects may also account for cases where the total topological charge is slightly less than the theoretical expectation of 12 (**Fig. 4a**).

We observe that 5-fold defects (mean radius of curvature at 453 Å) are located at more highly curved areas than the 6-fold hexamer (519 Å) and 7-fold defects (582 Å) (**Extended Data** Fig. 8a). Furthermore, we observe less regular defect arrangements with 4-fold or 8-fold valency at extreme local curvatures (**Extended Data** Fig. 7h,g), previously reported for lattices with weak interactions and may reflect local disorder of the HEF surface lattice^39^. Isolated 5-fold disclinations with only hexamers as adjacent oligomers, are observed only in smaller virus particles (ranging from 356 Å to 423 Å in radius) or large particles where local curvature is sufficiently high (mean radius of curvature at 420 Å). To further assess the impact of variable Gaussian curvature on defect formation^40–42^, we partitioned ellipsoidal virus particles into flatter top and bottom regions and more curved equatorial regions. Defects were present in both regions, although the fraction of defects was slighter higher in regions of greater curvature (**Extended Data** Fig. 8b).

The influenza C virus membranes generally display positive curvatures, with only occasional negatively curved regions. However, negatively curved regions, such as the invaginated region in the bi-lobed particle in **Extended Data** Fig. 9a, are denuded of glycoprotein, suggesting that HEF lattice contacts are incompatible with extended areas of negative curvature. Consequently, isolated 7-fold disclinations that could potentially occur on negatively curved surfaces were not observed, though defect pairs are found on the edge of the region.

Filamentous virus particles contain regions of cylindrical membrane where Gaussian curvature is close to 0. In **Extended Data** Fig. 9b, the matrix layer inside the membrane is a prominent feature and likely imparts the cylindrical shape of the particle as for other influenza viruses^43^. In theory, cylindrical crystals can consist of helical arrangements for specific values of radius or orientation of a hexagonal lattice, and this is approximately true for some highly ordered influenza C virus specimens^1,3^. However, in this example, grain boundaries are still observed in the hexagonal HEF lattice wrapping a cylinder (**Extended Data** Fig. 9b). Dislocation defects alongside vacant areas are associated with local bending or kinks that accompany slight reorientation of the helical axis, consistent with other studies on cylindrical crystals^44–46^. These may be associated with local variations in Gaussian curvature and a large principal curvature value in the direction orthogonal to the helical axis.

## Summary and significance

We studied the organisation of influenza C virus’s hexagonal HEF glycoprotein lattice, showing that physical principles of spherical crystallography govern its assembly. The system size and soft interactions of influenza C virus, make the growth of scars favourable compared to isolated disclinations, so that the number of excess defects scale with system size, consistent with theory. Direct imaging thus reveals new types of topological defects in assemblies of biological macromolecules and extends application of continuum elastic theory of lattice defects on curved surfaces at the nanometre scale.

Analysis of topological defects following in situ lattice reconstruction by cryo-ET and STA, as presented here, can be applied to other enveloped viruses^15^ and soft biological assemblies such as clathrin-coated vesicles^47,48^ and bacterial surface layers^49,50^ and may identify similar principles of assembly. Independent of a microscopic description of interactions, the main requirement for spherical assembly is that inter-molecular contacts can adapt to curved surfaces and form the described types of oligomers. More generally, our findings on the HEF lattice may therefore guide the design of new self-assembling biomaterials^51^ in 2D that exploit topological defects. Extending structural principles observed in symmetric and rigid viruses^52^, our study shows how a single protein component with weak interactions can cooperatively self-assemble flexible membrane containers of variable size suitable to the transport of complex and diverse cargoes.

Our studies have investigated static structures of fully assembled virus particles. Topological defects may play essential roles in dynamic structural changes in the lattice during the influenza C virus life-cycle. The tendency to form scars of neutral topological charge may, through screening, facilitate introduction of the disclinations^53^ needed for the transition of flat 2D membrane assemblies into spherical caps during initial budding events at the cell surface^54^. The HEF lattice both promotes membrane curvature and accommodates curvature generated by other viral components. Additionally, topological defects presumably play a role in lattice disassembly which occurs at low pH^55^ when the virus membrane fuses with host cell membranes to enter the cell. Thus, HEF lattice assemblies may be suitable for pH-sensitive delivery of drugs or genes by membrane fusion to cells. The role of specific molecular interactions between HEF glycoproteins, the membrane, and the matrix layer can now be explored in the context of the more general principles of assembly described here.

## Methods

### Virus growth, purification and vitrification for cryo-EM

Influenza C/Johannesburg/1/66 strain was grown in MDCK cells as previously described^3^ or in embryonated hen’s eggs^56^. For egg grown virus, influenza C/Johannesburg/1/66 virus was grown in the allantoic cavity of 10 day old embryonated hens’ eggs at 33 °C for 48 h. The virus was then harvested and purified by sucrose gradient centrifugation (30-60%). The virus pellet was resuspended in Tris-buffered saline supplemented with CaCl_2_ (with a final concentration of 50mM pH8.0 Tris, 100mM NaCl, 10mM CaCl_2_).

The purified virus was mixed with 10 nm Protein A gold particles (BBI) for use as fiducial markers. Quantifoil**^®^** R2/2 or R2/4 holey carbon grids with a 200 or 400 mesh were glow-discharged in the presence of amylamine. 4 μL of sample was applied to grids and then plunge-frozen using a Vitrobot™ (Mark III).

### Cryo-EM data collection

Tilt series for tomographic reconstruction and subtomogram averaging were collected using the Tomography software (Thermo Fisher) on a Titan Krios TEM (Thermo Fisher) equipped either with Falcon III detector operated in counting mode (4 frames/tilt angle) or with a post-column GIF Quantum energy filter (slit width of ± 10 eV) (Gatan) and a K2 summit direct electron detector (Gatan) operated in counting mode (4 frames/tilt angle). Three different tomography datasets of the purified virus have been collected under similar conditions (specific conditions are given in **Extended Data Table 1**) and used for further analysis. Dataset 1 was recorded as a bi-directional tilt series from −54° to +54° in 3° increments with a magnification corresponding to 2.23 Å/pixel (Falcon 3) and a dose of 1.36 e^-^/Å^2^ per tilt angle. Dataset 2, described in the previous article^3^ and re-processed and analysed as described below, was recorded as a bi-directional tilt series from −54° to +54° or −42° to +54° in 3° increments with a magnification corresponding to a 2.20 Å/pixel (K2) and a dose of 1.57 e^-^/Å^2^ per tilt angle. Dataset 3 was a bi-directional tilt series, from 0 to ± 60° or −42° to +54° in 3° increments with a magnification corresponding to 1.38 Å/pixel (K2) and a dose of 2.10 e^-^/Å^2^ per tilt angle.

### Image processing and subtomogram averaging

All three datasets were pre-processed in the same way as previously described^3^. Movie frames were motion corrected and dose weighted using Alignframes from the IMOD package^57^. Tilt series were aligned using Etomo from the IMOD package^57^. The contrast transfer function (CTF) was estimated using CTFFIND4^58^ and tomograms were CTF corrected and reconstructed using novaCTF^59^. For visualisation purposes, the tomograms used in **Fig. 1b-d** were treated with Topaz^60^ for denoising and IsoNet^61^ to reduce effects of the missing wedge of data due to the limited tilt range of the specimen stage.

For subtomogram averaging, the three datasets were processed separately with similar procedures (**Extended Data** Fig. 1). Initial subtomogram coordinates were seeded and picked using Dynamo^62^ membrane and vesicle models from SIRT-filtered tomograms (IMOD) to identify the positions and orientations of HEF trimers based on the geometry of each virus particle. These coordinates were picked with a spacing interval of 30 Å which is less than the inter-dimer distance between two HEF molecules (∼85 Å), to ensure all HEF molecules were picked.

Subtomograms were extracted from the original tomogram reconstructed by weighted back projection (WBP), using 4× down-sampling to aid computational efficiency, with a box size of 64^3^ pixels in RELION-3.0^63,64^. A small subset of subtomograms (∼15K) was used to generate an initial reference map for downstream processing. Two rounds of 3D classifications in C1 symmetry were performed to get a homogeneous set of subtomograms (∼86K), after removing the duplicates from over-sampled picking. These subtomograms were re-extracted from un-binned tomograms (256^3^) and subjected to refinement with C3 symmetry. The refined map was post-processed with a soft mask (**Fig. 2a** and **Extended Data** Fig. 10a).

For reconstructions of oligomers, individual HEFs (4× downsampled) were symmetry expanded in C3 in RELION followed by a shift to create new coordinates at any one of the three nearest oligomer centres. 3D classifications in C1 identified hexamers (∼35K), pentamers (∼3K), and heptamers (306). These classes were re-extracted from un-binned tomograms and refined with C6, C5, and C7 symmetry, respectively (**Fig. 2b-d** and **Extended Data** Fig. 10b**-d**), and the symmetrised maps overall match with the 3D classes determined in C1.

At bin4, subtomograms of pentamers and heptamers were symmetry-expanded in C5 and C7, respectively, and shifted to re-centre on the dislocation pair dimer interface. Approximately 13,000 particles were obtained via enlarged box extraction (∼706 Å), potentially including duplicates. Following duplicate removal, two rounds of 3D classification, using a mask focused on the “5-7” defect densities, yielded 588 particles. Subsequent refinement and post-processing produced a 27.2 Å map that consists of a “5-7” dislocation pair and a hexamer of HEF trimers, at 4× down-sampling (**Fig. 2e** and **Extended Data** Fig. 10e).

Models for each oligomer map (**Fig. 2b-d**) and the composite oligomer map (**Fig. 2e**) were built through rigid fitting of individual HEF trimers (PDB: 6YI5) into the STA maps using UCSF ChimeraX^65^. Interface residues and buried surface area were calculated using PISA^66^ (**Extended Data** Fig. 2a).

### Template-matching generated surface lattice models

We performed template matching using STA maps for pentameric, hexameric, and heptameric oligomers, as well as individual HEF trimers against the weighted back-projection (WBP)-reconstructed tomograms (4× downsampled) using pytom-match-pick^67^.

Cross-correlation (cc) peaks (threshold of 0.15 and a particle diameter of 170 Å for peak detection) form regularly spaced 3D points (**Extended Data** Fig. 3a), consistent with its arrangement as a hexagonal surface lattice and provide isotropic coverage across the virus surface. The oligomeric type was assigned as the highest scoring template within a 100 Å search radius.

Hexamer centres were used to identify HEF trimer positions. These were obtained by C6 symmetry expansion of hexamers followed by a positional shift to HEF trimer vertices (**Extended Data** Fig. 3b). This step locates trimers on both hexamers and defect region, as pentamers and heptamers are typically adjacent to multiple hexamers in the context of a lattice. Unique HEF trimer positions were chosen as the trimer with the highest cc score within a 60 Å distance.

Coordinates of the single HEF trimer vertices and oligomer centres of HEF trimers (**Extended Data** Fig. 3c) were assigned to individual virus particles based on proximity to the virus particle models generated using Dynamo’s particle picking (**Extended Data** Fig. 3d). 3D surface meshes were reconstructed for each virus particle via *trimesh* in Python, containing both HEF trimer vertices and oligomer centres. These virus surface meshes exhibit gaps due to incomplete surface coverage across the virus surface. To fill in gaps and produce a continuous and closed surface, we implemented a radial basis function interpolation^68^ to construct an implicit surface, followed by *trimesh* Taubin smoothing^69^. These complete surface models enabled identification and exclusion of out-of-plane lattice positions. HEF trimers and oligomer centres were removed if the distance from the fitted surface exceeded 30 Å. Surface lattice coverage was then quantified as the ratio of the observed glycoprotein mesh area to the total interpolated surface area.

### Curation of highly covered surface lattice models

Many virus particles exhibited a clustered radius distribution near ∼50 nm. Models with high surface coverage were selected across a range of particle sizes. Models that have a surface coverage greater than 80% are selected for curation. In addition, for models outside the 50 nm radius cluster, a threshold of >60% was used to ensure the diversity of particle size and shape in downstream analyses.

A total of 42 virus particles were manually curated (**Extended Data Table 2**). When modelling the surface lattice, the built models were visualised using ArtiaX^70^, which renders individual surface glycoprotein trimers as volumetric densities and centres of oligomeric assemblies as spherical markers, coloured in red, blue and yellow representing pentamers, hexamers and heptamers, a default representation across all figures.

We performed manual corrections on these curated models to ensure accurate placement of HEF trimer vertices and oligomeric centres through an interactive procedure in ChimeraX and ArtiaX (**Extended Data** Fig. 4a,b). Oligomer types were reassigned when necessary based on the number of surrounding HEF trimers (mostly 5, 6, or 7-fold, and, in rare cases, 4-fold or 8-fold), with the expected lattice spacing (∼85 Å). Missed oligomer markers were added at the geometric centroids of the HEF trimer polygons, while existing oligomers lacking sufficient surrounding HEF trimers were removed. Concurrently, we completed partial polygons by adding HEF trimers when three plausible neighbour interactions can be found. HEF trimers exhibiting severe clashes or improper orientations were either reoriented or removed. The final curated lattice models, comprising single HEF trimers, and centres of pentamer, hexamer, and heptamer of HEF trimers, were saved in RELION .star format for downstream analysis.

The accuracy of oligomer type assignment was assessed by counting the number of surrounding trimer vertices. For each oligomer centre, coordinating trimer vertices were identified within an expanded (1.4×) radius of the expected distance for regular polygons (101 Å for pentamers, 119 Å for hexamers, and 138 Å for heptamers). This confirmed high-confidence oligomer type assignment (700/702 for pentamer, 3797/3829 hexamer, 382/414 for heptamer). To quantify geometric regularity of oligomers, trimer vertices were projected from 3D space onto a 2D plane defined by the first two principal components (PC1 and PC2) using principal component analysis (PCA) using the *scikit-learn*^71^ package in Python, which captures the best-fit planar orientation. The projected vertices were then aligned to the corresponding ideal regular polygon using the Kabsch algorithm^72^, and root-mean-square deviation (RMSD) values for HEF trimers were calculated between observed and ideal positions, with hexamers being the most regular (6.5 Å), followed by pentamers (6.8 Å) and heptamers (10.1 Å).

To assess the surface protein modelling quality of each virus particle, steric clashes between modelled glycoproteins were assessed via back-plotting atomic models for each glycoprotein trimer (PDB: 6YI5) in the tomogram space. A volume overlap percentage (a concept analogous to molecular packing function^73,74^ used for measuring steric overlaps) was then calculated based on a simulated molecular envelope generated from the atomic model using the ChimeraX *molmap* tool at 10 Å resolution, with volume calculated at a density threshold of 0.10, a volume threshold sufficiently covering the model. An ideal glycoprotein packing would yield a 0% overlap. Across the 42 built virus surface models, the average overlap was 0.86% of the modelled surface lattice glycoprotein volume, indicating good spatial compatibility and minimal steric clashes between neighbouring surface glycoproteins.

To further confirm the modelling of oligomers, the final oligomeric positions were reimported into RELION and re-extracted as subtomograms at bin4. Using Euler angles derived from template matching, subtomograms labelled as 5, 6, and 7-fold were able to be reconstructed into their expected pentameric, hexameric, and heptameric assemblies at C1 (**Extended Data** Fig. 5a). After further refinement in C1, masking and post-processing, the resulting reconstructions revealed oligomer structures consistent with those obtained from previous STA classifications. Back-plotting confirmed the consistency of the oligomers with the modelled lattice (**Extended Data** Fig. 5b). The agreement between these two structural determination approaches supports the overall accuracy of oligomer type assignment, as well as their positioning and orientation obtained from the modelled surface lattices.

Additionally, to evaluate sensitivity of the procedures to the missing data wedge, we partitioned the oligomers based on their orientations and performed subtomogram averaging for both orientations, using reference-free alignment at C1 symmetry, without masking: (1) top (0° < θ < 40°) and bottom (140° < θ < 180°) views (**Extended Data** Fig. 6a), and (2) side views (75° < θ < 105°) (**Extended Data** Fig. 6b), where *θ* is the assigned Euler orientation angle in RELION following the Euler angle convention described in Heymann et al.^75^. These refined oligomer maps were plotted back to their original positions based on their refined 3D coordinates and orientations, showing their geometric consistency that match with adjacent oligomers in both orientation groups (**Extended Data** Fig. 6).

### Topological and geometric analysis of the modelled lattice

HEF trimer vertices and oligomer centres were imported into PyMeshLab^76^ to reconstruct the lattice into a mesh and enable mesh-related computations. The curved lattice mesh was first triangulated using the ball-pivoting algorithm (BPA)^77^, followed by Laplacian smoothing^78^ and two iterations of Least Squares Subdivision Surfaces^79^ to generate an oversampled mesh.

Topological properties were quantified for each modelled surface lattice. Given the pleomorphic nature of the virus particles, including ellipsoidal morphologies, the effective (area-equivalent) radius *R* was derived from the surface area of the interpolated full mesh, with 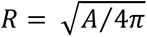, where *A* is the area of surface located at the layer of HEF trimer centroid. This radius was used to calculate the system size, defined as *R/a*, with the measured distance between two single HEFs (∼ 85 Å) as the spacing *a* of the system.

Local Gaussian curvature and principal radius of curvature, calculated as 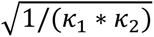, where *κ_1_* and *κ_2_* are the principal curvatures, were computed at all vertices of the oversampled mesh via scaled quadric surface fitting (see example in **Extended Data** Fig. 4d and **Extended Data** Fig. 8). These curvature values were subsequently mapped back to the original lattice points via nearest-neighbour assignment to the corresponding mesh vertices. To further verify the curvature estimation, we examined virus particles with distinct morphologies (e.g., ellipsoidal and filamentous virus particles). The resulting curvature distributions matched expected geometric features: high radii of curvature on flatter surfaces, lower values in the highly curved equatorial belt (**Extended Data** Fig. 8c), and near-uniformly high values along the cylindrical surface of filamentous virus particles, consistent with near-zero Gaussian curvature (**Extended Data** Fig. 9b). Shape index in **Extended Data** Fig. 9a is defined as in Koenderink and van Doorn^80^.

### Plotting and Statistical analysis

The quantitative measurements of each modelled virus surface lattice are provided in **Extended Data Table 2**.

Surface lattice coverage, calculated as the ratio of the modelled glycoprotein mesh area to the total interpolated surface area, was used to extrapolate the total topological charge. For each virus lattice model, the number of oligomers were extrapolated for figure plotting in **Fig. 4a,c**, via surface coverage for the total number if the lattices were fully covered manifolds. The number of excess defect pairs, calculated as (*n_5_* + *n_7_* - 12)/2, was then divided by 12 (a full icosahedron would have 12 5-fold disclinations) to obtain the number of excess defect pairs per disclination, a metric previously used in the analysis of colloidal spherical crystals^5^.

Using the SciPy^81^ package in Python, a linear least-squares regression was performed to determine the slope of 0.56 and the Pearson correlation of 0.78 (*p* = 1.53 × 10^-9^), incorporating virus particles both with and without matrix layers. For particles possessing a matrix layer, the system size *R/a* clustered around a mean of 6.06, with a standard deviation of 0.30 of excess defect pairs. A linear regression plot of the excess number of defect pairs and the system size *R/a* was created using the Seaborn^82^ plotting package (**Fig. 4**).

Grain boundaries (or scars) were identified as HEF trimers forming the vertices of pentamer and heptamer defects (examples highlighted in orange in the 3D back plots in **Fig. 1e-g** and **Fig. 3**). For each reconstructed surface lattice model, the total number of grain boundaries and their corresponding net topological charge were enumerated and are summarised in **Extended Data Table 2**. Scar length (**Fig. 4b**) was quantified as the angle between vectors from the virus particle centre to the oligomer centres marking the start and end of the scar. The distribution of scar lengths was visualised as a histogram using Seaborn.

Virus particles were grouped into spherical and ellipsoidal types by aspect ratio (semi-major axis divided by semi-minor axis, threshold = 1.2). For spherical virus particles, curvature radii of all oligomer centres were grouped by oligomer type and pooled across all viewing orientations. Kernel density estimate (KDE) plots describe curvature distributions for pentamers, hexamers, and heptamers (**Extended Data** Fig. 8a). For ellipsoidal virus particles, oligomers were further separated into top/bottom views (0–40° and 140–180°) and side views (40–140°). In these particles, all three oligomer types exhibited higher mean radii in the top/bottom views compared with the side views, consistent with their short axis normal to the plane of the ice film. In ellipsoidal virus particles, a defect fraction number, which measures the number of defects divided by total morphological units, were calculated between top/bottom and side regions (**Extended Data** Fig. 8b).

## Extended Data

**Extended Data Fig. 1.**
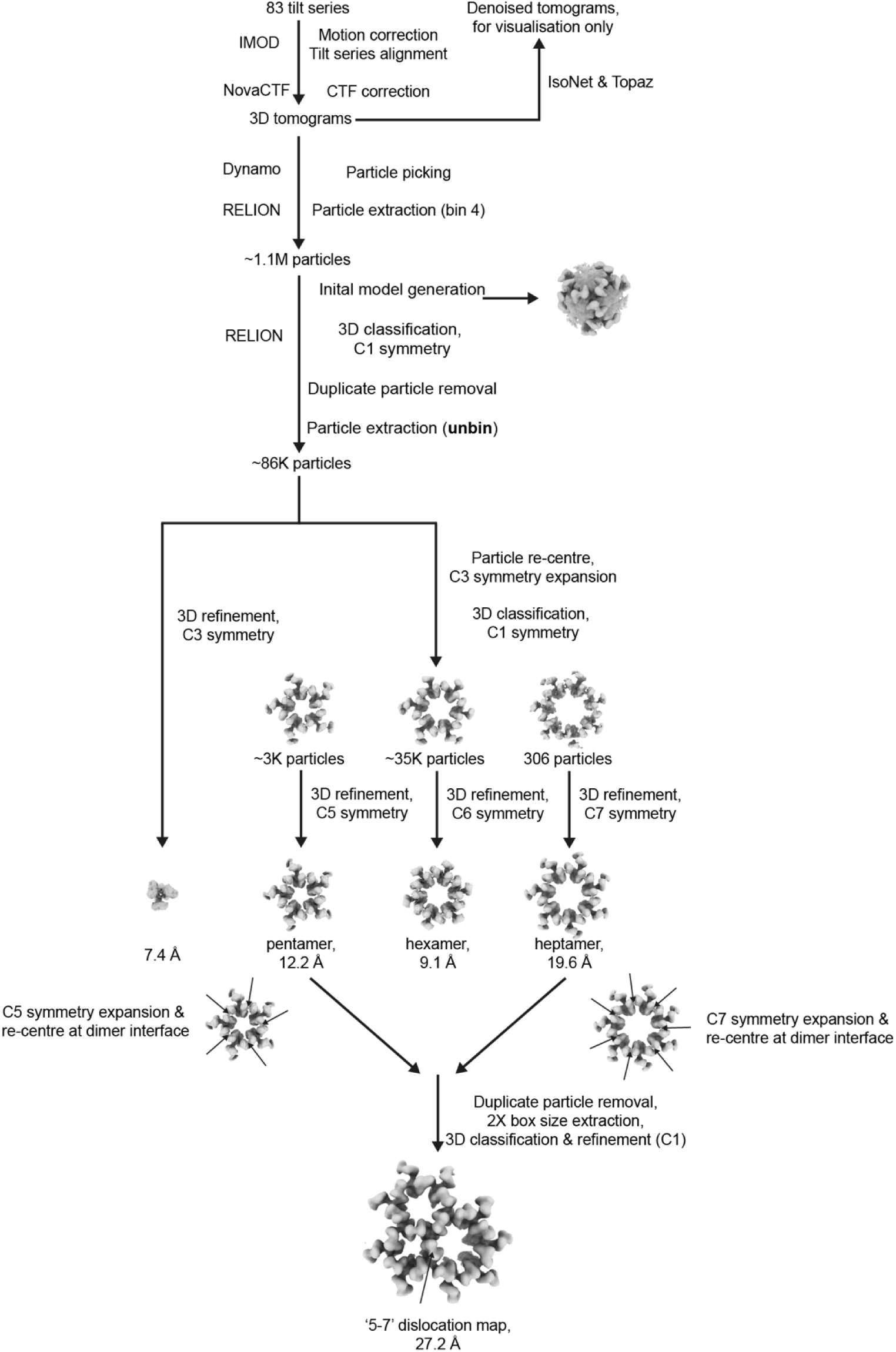
The computational workflow for tomogram reconstruction and subtomogram averaging (STA). The five maps from Fig. 2a**-e** were obtained using this processing workflow as described in **Methods**, reconstructed with C3, C6, C5, C7 and C1 symmetry, respectively.

**Extended Data Fig. 2.**
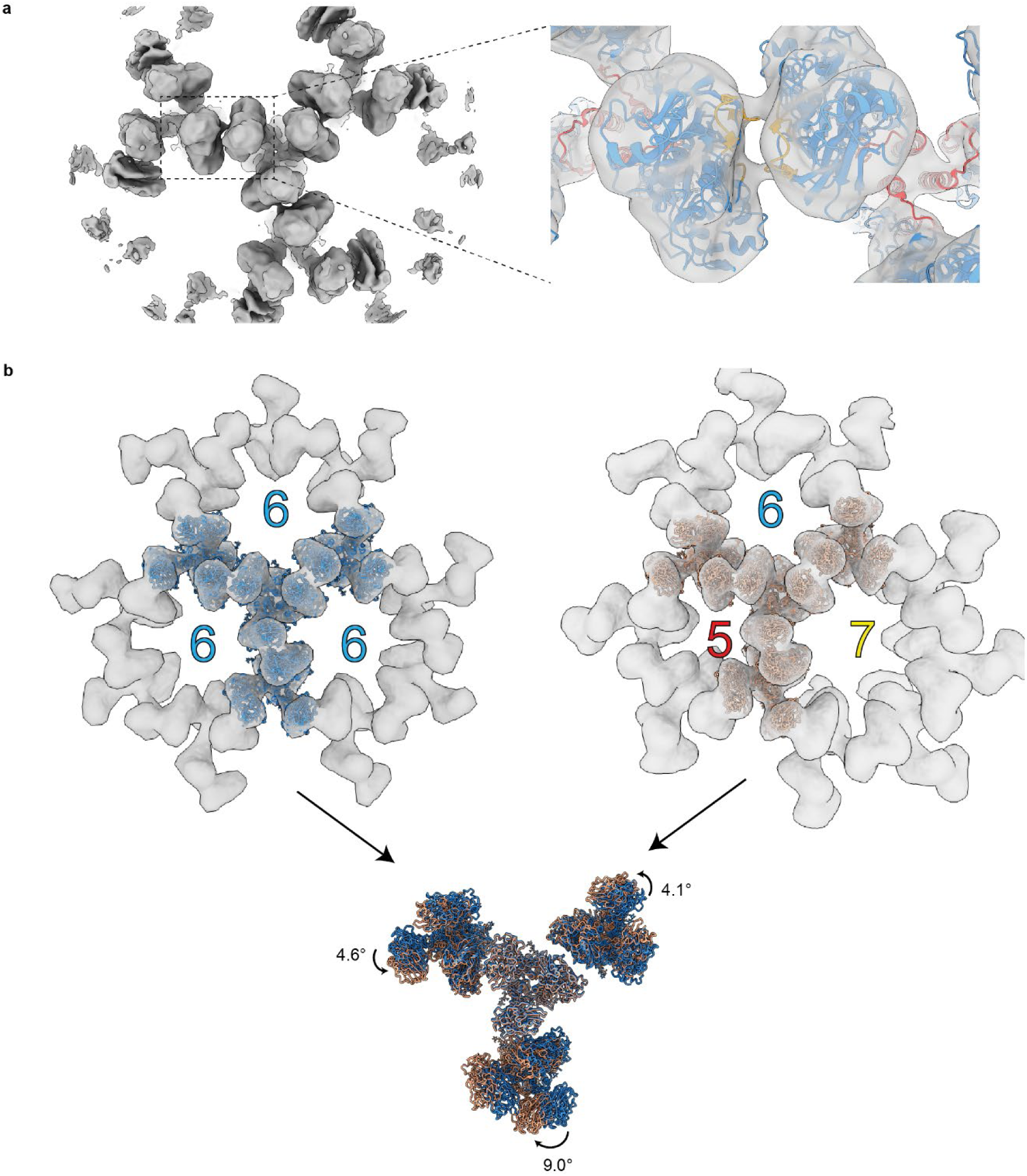
Molecular model of the dimer interface of HEF trimers and structural comparison of the oligomeric assemblies. **a**, Two HEF trimers (PDB: 6YI5) docked into the full STA reconstruction from Fig. 2a without central trimer mask applied, showing density for three neighbouring HEF trimers interacting with the central trimer. A magnified view of the dimer interface highlights two surface loops (orange) between the HEF1 membrane-distal domains of adjacent monomers, with an interface of buried surface area 209.3 Å², contributing to approximately 2.03 kcal/mol. **b**, Comparison of HEF trimer configurations within a pure hexameric lattice (STA map from Fig. 2a, with 4 x down-sampling and a larger box size) and a 5–7 dislocation (map from Fig. 2e), with four trimer models fitted into each map. Trimers in the pure hexameric lattice are coloured blue, and those at dislocations are coloured coral. One central trimer is shared by three neighbouring oligomers. Comparing the two cases, the neighbouring trimers show the following rotations with respect to the central trimer: 4.6° for the transition from “6–6” to “6–5”, 4.1° for “6–6” to “6–7”, and 9.0° for “6–6” to “5–7”. Angles were measured between the centroid of the central trimer and those of the neighbouring trimers, indicating the conformational adjustments required for defect insertion in the lattice.

**Extended Data Fig. 3.**
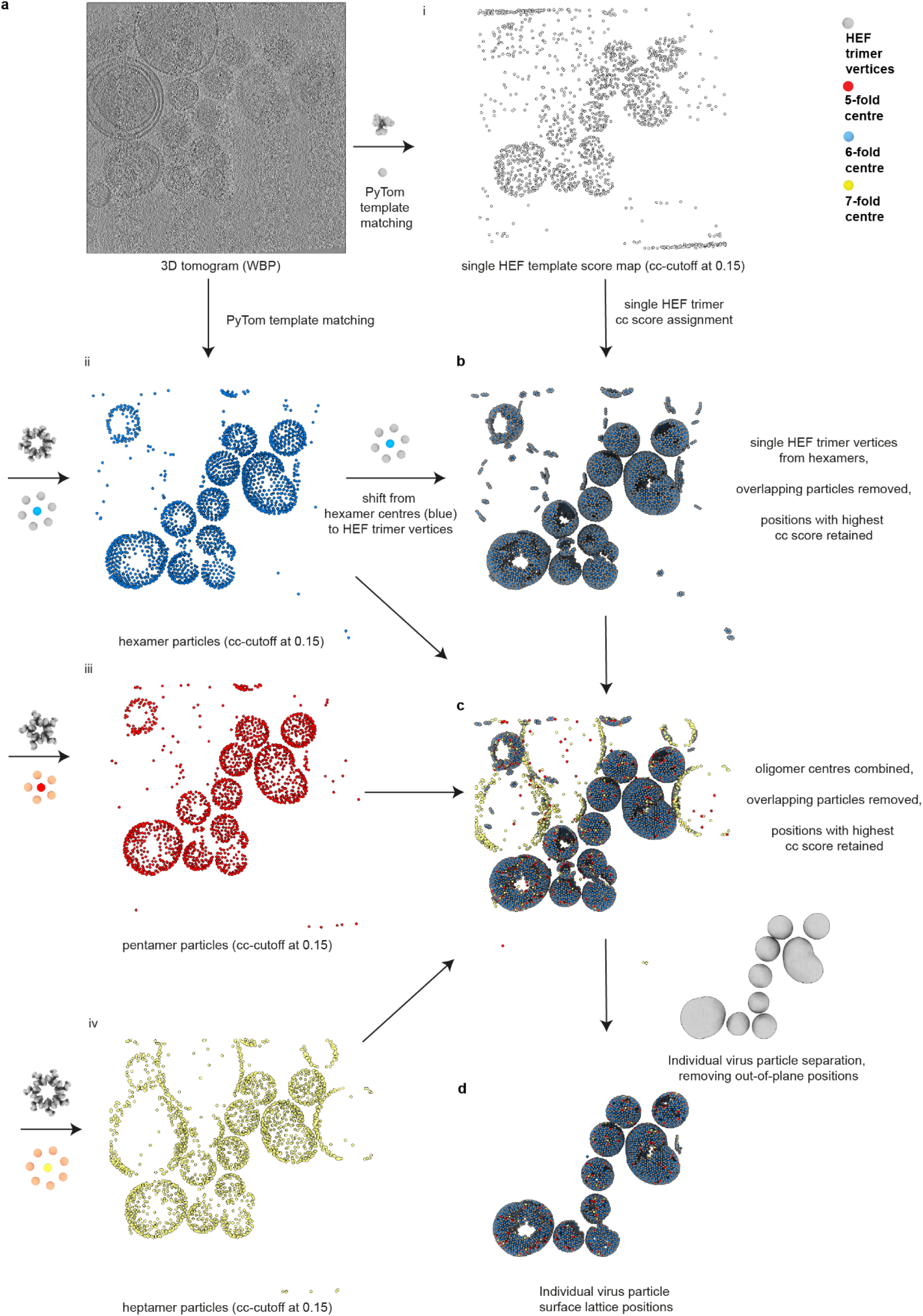
Automated surface lattice modelling using template matching. **a,** For each WBP-reconstructed tomogram (at bin 4), four map templates (single HEF trimer, and oligomers comprising pentamers, hexamers, and heptamers of HEF trimers) were used to identify putative surface lattice positions. Cross-correlation (cc) maps were generated voxel-wise for each template, yielding particle coordinates for (i) individual trimer positions (grey), (ii) hexamer centres (blue), (iii) pentamer centres (red), and (iv) heptamer centres (yellow), using a cc threshold of 0.15. **b,** As individual trimer positions in panel **a**i from template matching exhibited low signal-to-noise ratio, lattice trimer vertices were instead modelled using the vertices of hexamer centres. Duplicate vertices were removed within a 60 Å search radius, retaining the highest-ranking cc match, with cc scores taken from the individual trimer cc map (panel **a**i). **c,** The three sets of oligomer centres were combined and duplicates removed within a 100 Å search radius, retaining the highest-ranking cc match. The retained centre and its highest cc score assigned the initial oligomer type. **d,** Surface lattice positions were then grouped by virus particle identity based on initial surface-model particle picking in Dynamo and filtered using surface morphology to remove off-surface particles.

**Extended Data Fig. 4.**
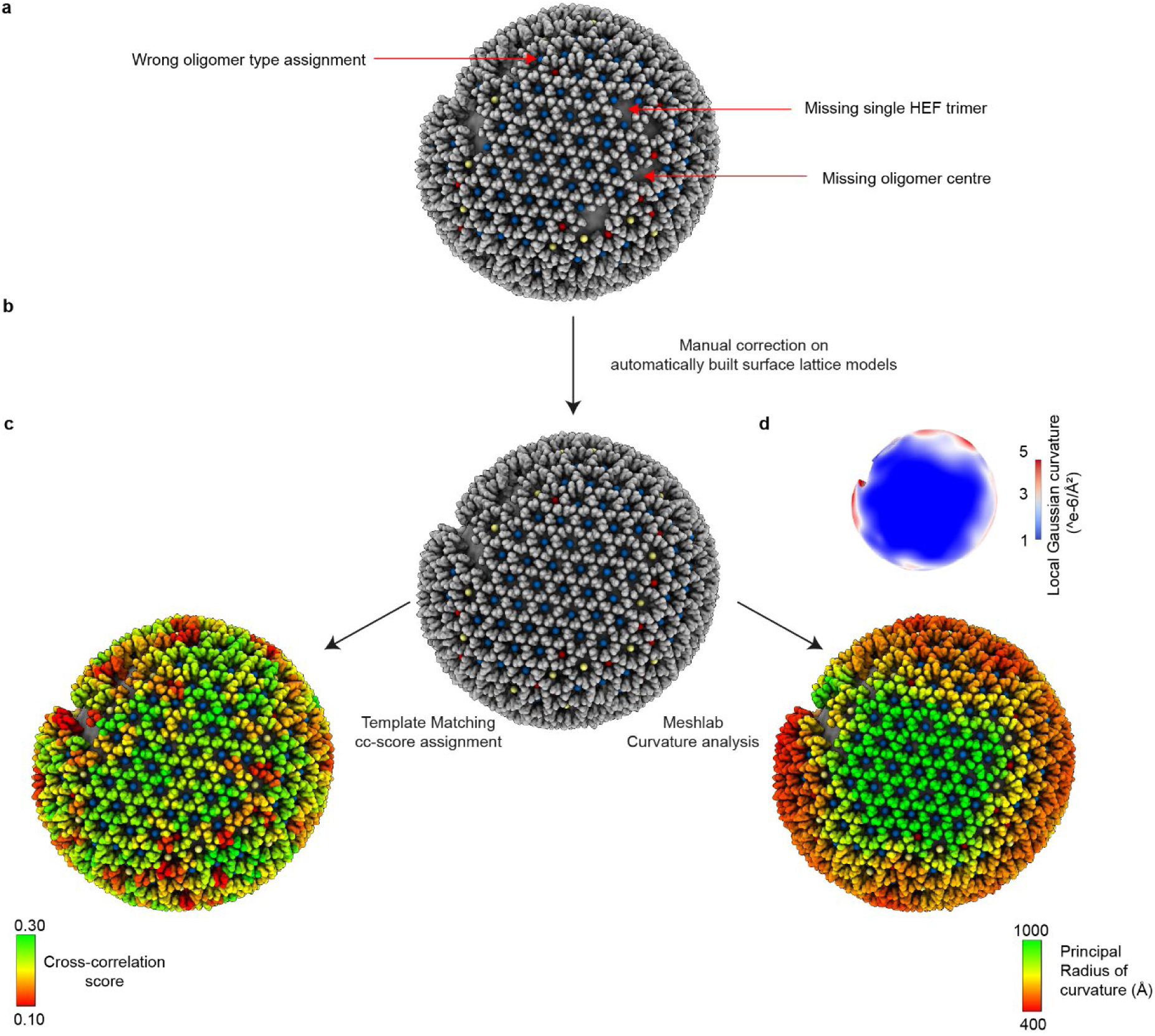
Curation and analysis of surface lattice models. **a,** Example of an initial surface lattice model (The same virus particle is shown in Fig.1) generated by automated template matching, which contains false-positive and false-negative lattice positions, misoriented trimers and incorrectly assigned oligomer types. **b,** To ensure accurate modelling of the surface lattice, the HEF trimer positions, orientations and the oligomer centre assignments were manually corrected to match the local geometric environment. **c,** To evaluate local trimer modelling quality, cross-correlation scores (0.10 to 0.30) from single-trimer template matching were assigned to each lattice vertex and visualised by colour. **d,** The corrected surface lattice was converted into an oversampled triangulated mesh using MeshLab for local Gaussian curvature calculation. An example ellipsoidal virus particle is shown coloured by radius of curvature (400–1,000 Å).

**Extended Data Fig. 5.**
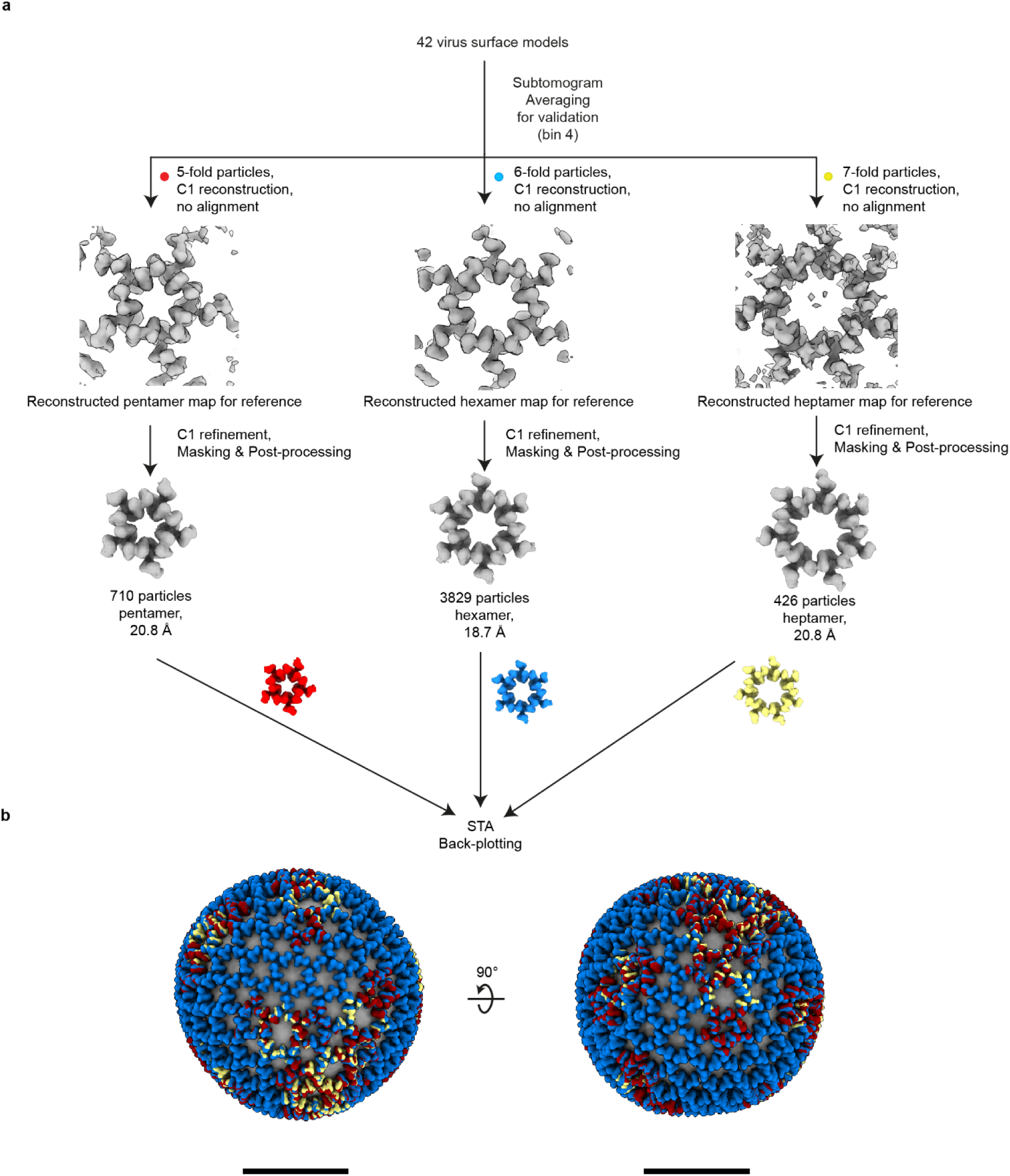
Subtomogram averaging (STA) and back-plotting of modelled oligomers to confirm oligomer type assignment. **a,** STA of oligomers was performed based on assigned type and position. C1 reconstructions (bin 4) of the three types of oligomers resulted in the expected polygonal structure (zoomed in on the central portion of the STA map). STA refinements in C1 produced maps with the corresponding densities of oligomeric assemblies of HEF trimers, which are consistent with those obtained via STA 3D classification (see **Fig.2b-d**). **b**, These maps were then back-plotted onto their original positions within the tomograms using the 3D orientations and coordinates determined by STA. The resulting overlay of three oligomer back-plot shows that the HEF trimer vertices from each oligomer align well with neighbouring oligomers, exhibiting geometric consistency without obvious structural clashes. The example shown here is from virus particle no. 29 in **Extended Data Table 2**. Scale bar: 50 nm.

**Extended Data Fig. 6.**
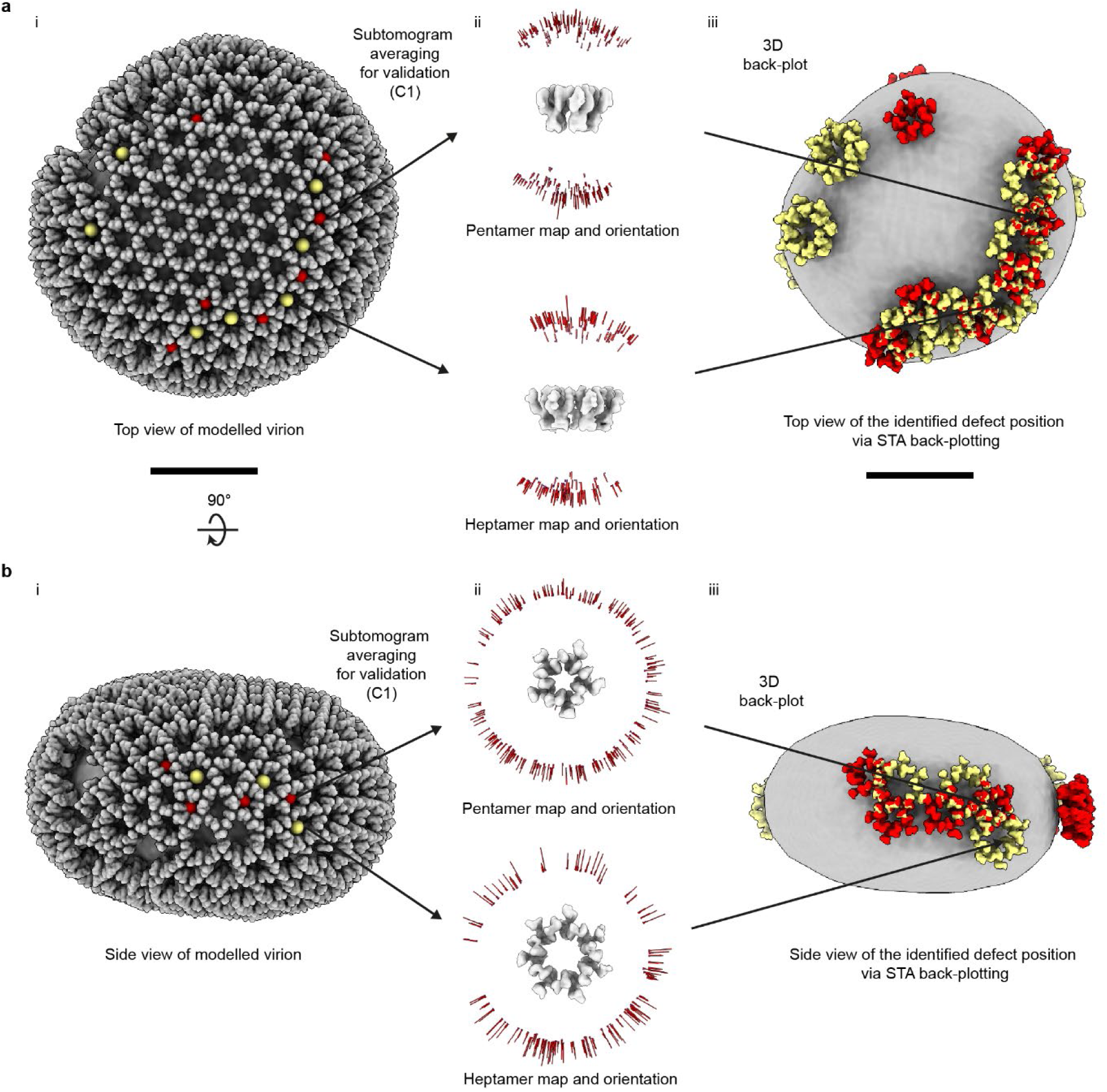
Subtomogram averaging (STA) of modelled defects across different viewing orientations. The defect oligomers were correctly identified and aligned in two distinct viewing orientations of the tomogram, contributing to isotropic surface models. The example workflow demonstrates (i) the modelled virus particle surface, (ii) the STA maps of the corresponding defect oligomer and the orientation distribution (shown in red bars) and (iii) back-plotted STA structures in 3D (5-fold in red and 7-fold in yellow). The identified defect oligomers were extracted from modelled virus particles, separated into two different projection views for STA validation: **a,** top and bottom views (subtomogram tilt angle 0° < θ < 40° and 140° < θ < 180°); **b,** side views (75° < θ < 105°). The same virus particle is shown in Fig.1. Scale bar: 50 nm.

**Extended Data Fig. 7.**
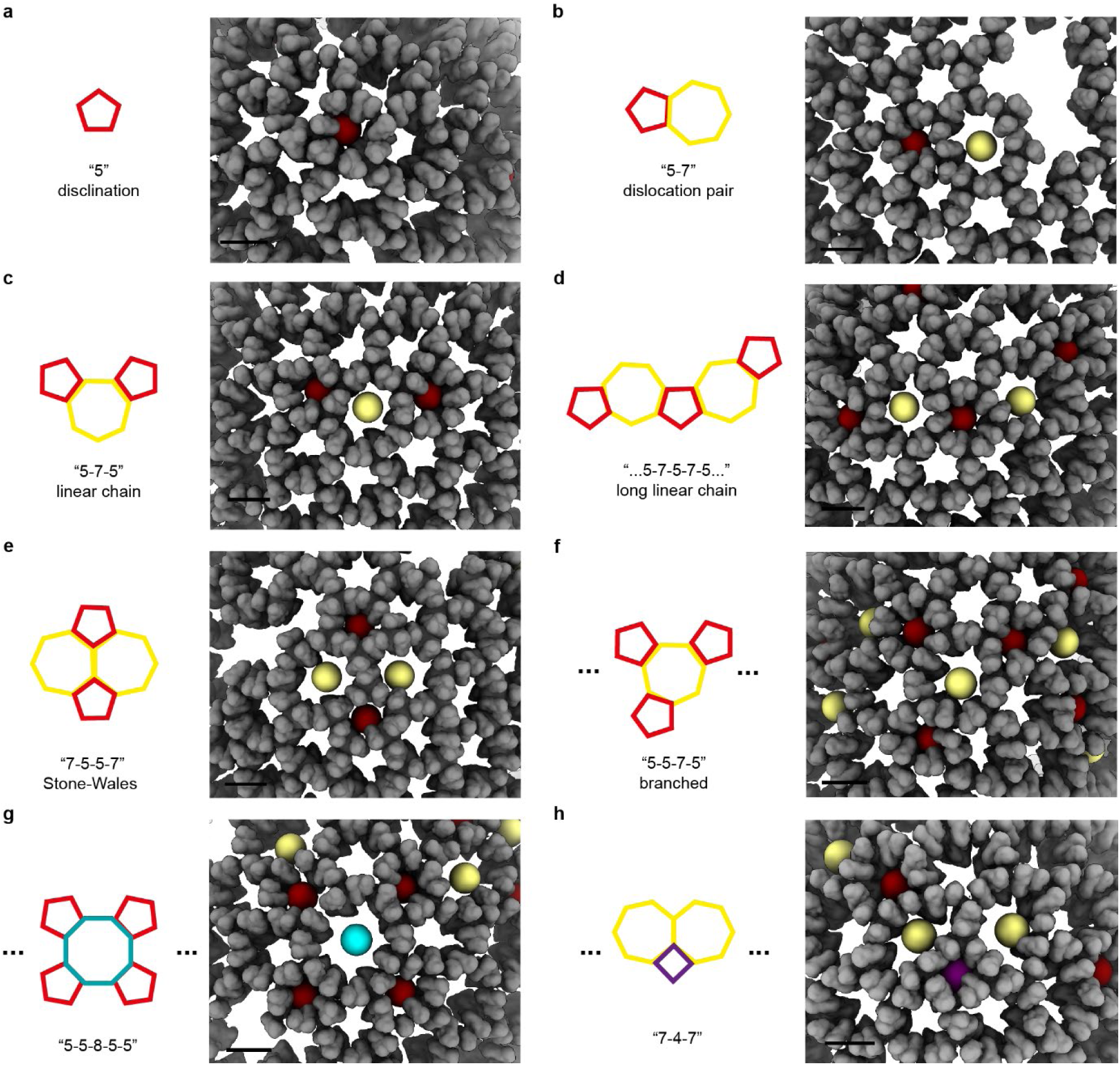
Point defect types observed in the modelled lattice volume (*n*-fold defect type carries a net topological charge q = 6 - *n*). **a**, Isolated 5-fold disclination, surrounded by five hexamers. **b**, “5-7” defect pair or dislocation, adjacent to a void region. **c**, “5-7-5” linear dislocation chain. **d**, Long dislocation chain consists of more than one excess dislocation pair (…“5-7-5-7-5”…). **e**, “7-5-5-7” Stone-Wales defect. **f**, Branched “5-5-7-5” dislocation. **g**, “5-5-8-5-5” dislocation. **h**, “7-4-7” dislocation. Scale bar: 10 nm.

**Extended Data Fig. 8.**
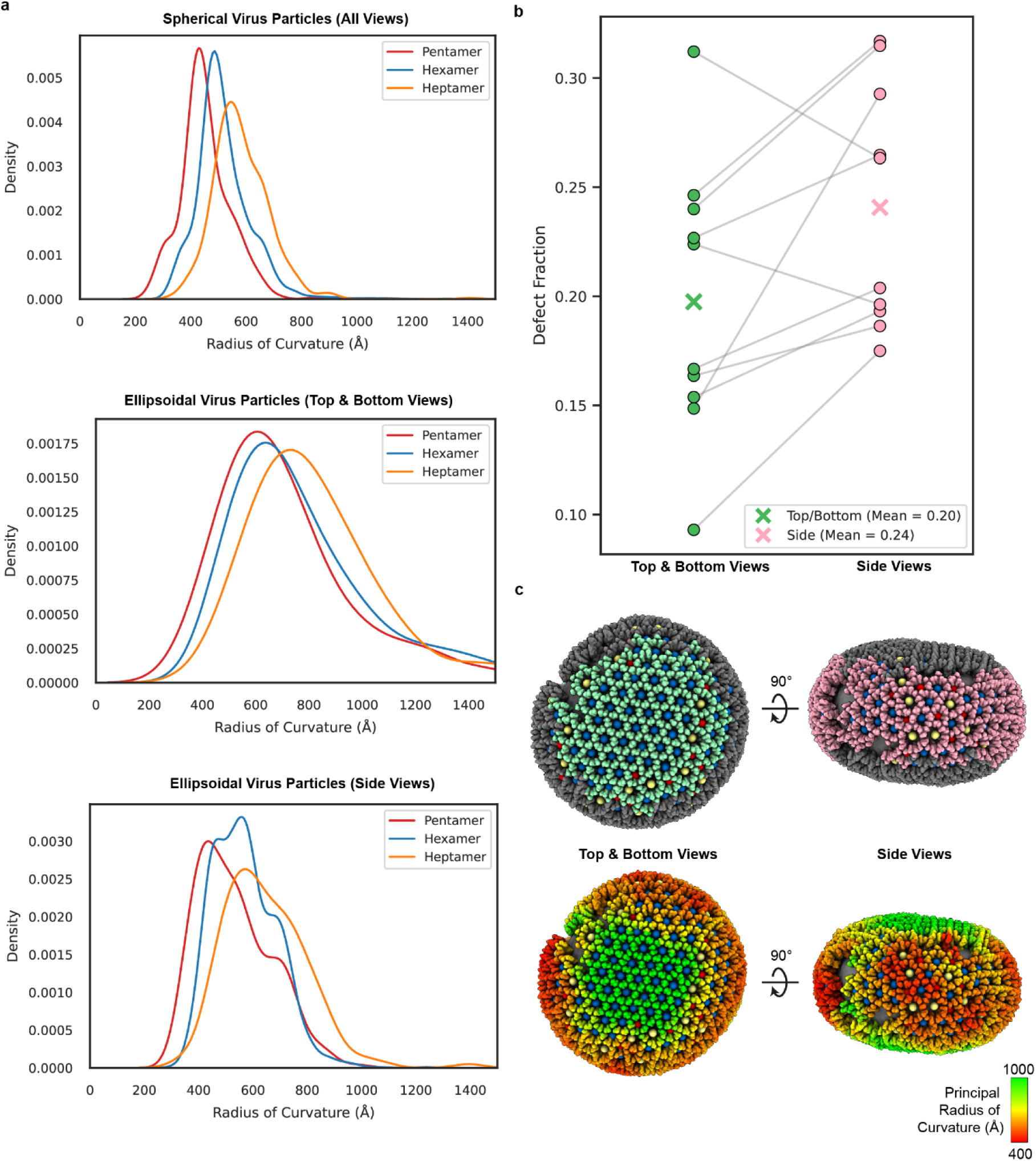
Distribution of topological defects in virus particles with varying morphology and local curvature. **a**, Kernel density estimates (KDEs) of oligomer distribution as a function of local radius of curvature (Å). For spherical particles (aspect ratio ≤ 1.2), the mean radius of curvature was smaller for pentamers (453 Å, n = 749) than for hexamers (519 Å, *n* = 4122) and heptamers (582 Å, *n* = 431). For ellipsoidal particles (aspect ratio > 1.2), oligomers were analysed separately in top and bottom views (tilt 0–40° and 140–180°) and in side views (tilt 40–140°). In top and bottom views, pentamers (727 Å, *n* = 108), hexamers (819 Å, *n* = 765), and heptamers (850 Å, *n* = 87) all exhibited larger radii compared with side views, where pentamers (513 Å, *n* = 89), hexamers (562 Å, *n* = 452), and heptamers (637 Å, *n* = 61) displayed smaller radii. **b**, The defect fraction for different top and bottom (green) and side (pink) views of ellipsoidal virus particles (*n* = 10, aspect ratio > 1.2). Defect fraction, computed as number of defects divided by the total number of morphological units in the area, was higher in the more curved side regions (0.24) than in the flatter top/bottom regions (0.20) (*t*-test, *p* = 0.035). **c**, Surface lattice model of an ellipsoidal virus particle, coloured according to regions defined in panel **b** (green, flatter top/bottom; pink, more curved side belt). Lower panels show the corresponding surface model, with radius of curvature values colour-coded from 400 Å (red) to 1,000 Å (green). Oligomer centres are denoted by coloured spheres: red (pentamers), blue (hexamers), and yellow (heptamers). The same virus particle is shown in Fig.1.

**Extended Data Fig. 9.**
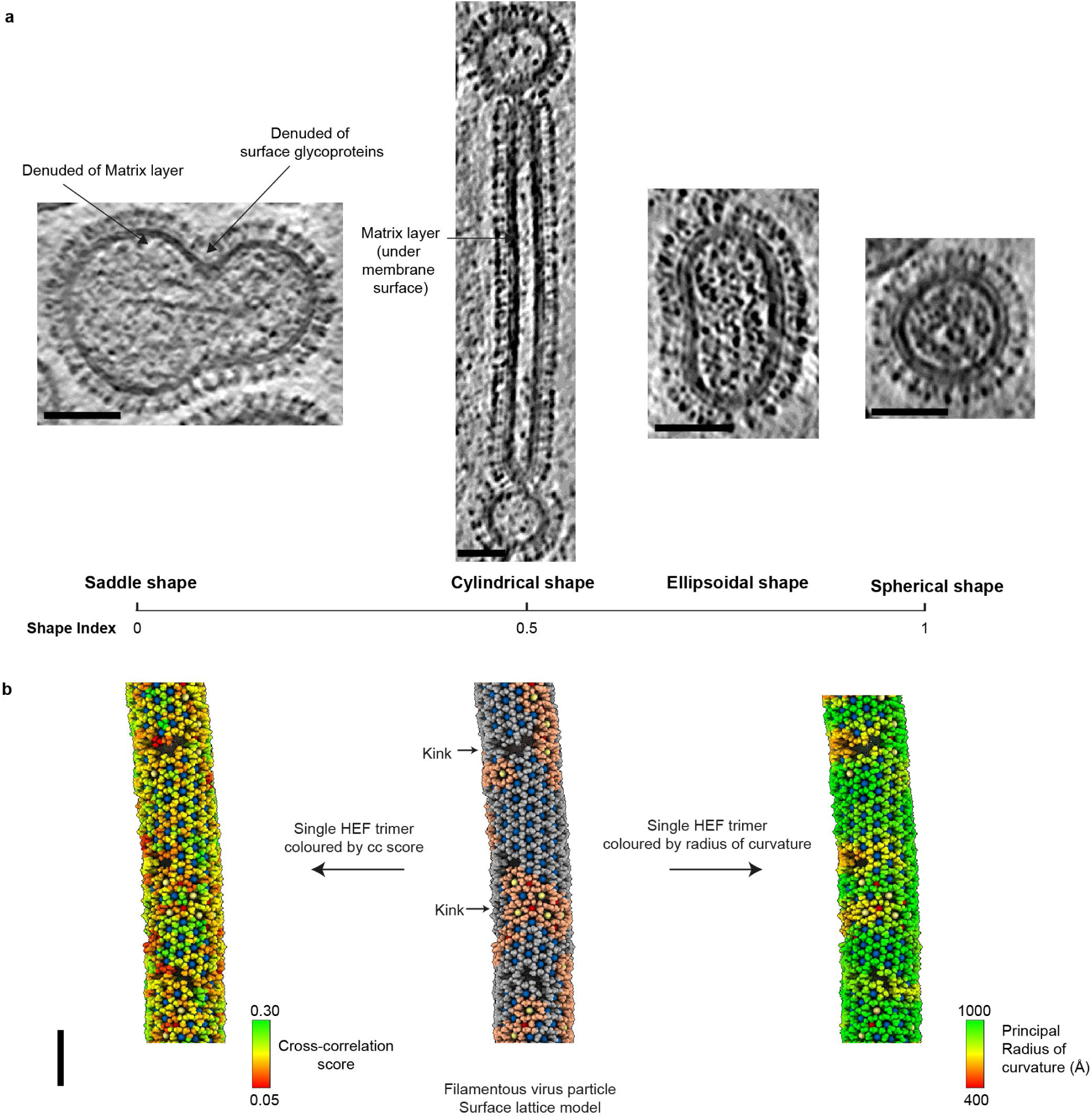
Grain boundary defects in HEF lattices of non-spheroidal and spheroidal virus particles. **a,** Representative virus particles are arranged from left to right by increasing shape index of local regions, calculated from the principal curvatures *κ_1_* and *κ_2_*. This arrangement covers a range of morphologies, from saddle shape (shape index of 0) to spherical shape (shape index of 1). Examples include: a bi-lobed virus particle with a local saddle-shaped region, featured by one positive and one negative principal curvature. This virus particle lacks a matrix layer and is largely denuded of surface glycoproteins in the negatively curved neck region; a virus particle with an extended cylindrical segment in the middle. The elongated filament segment exhibits one principal curvature near zero and the other high, with a matrix layer underlying the membrane; an ellipsoidal virus particle; and a nearly spherical virus particle. **b**, Surface lattice model of the cylindrical segment of a virus particle from panel **a** (∼370 nm in length and ∼30 nm in radius). The virus particle exhibits low Gaussian curvature across most of its surface, corresponding to nearly flat regions with large radii of curvature. Localised bending (kink) along the filament introduces curvature variation, which facilitates the formation of numerous lattice defects. HEF glycoproteins assemble into a predominantly hexagonal lattice, with an approximate helical organisation about the filament axis. HEF trimer positions are colour-coded to distinguish between regular hexagonal domains and grain boundary regions, following the same scheme as Fig.1. The surface model is also coloured by trimer positions according to cross-correlation scores (0.05 to 0.3) from template matching, reflecting the quality of individual position assignments. Scale bar: 50 nm.

**Extended Data Fig. 10.**
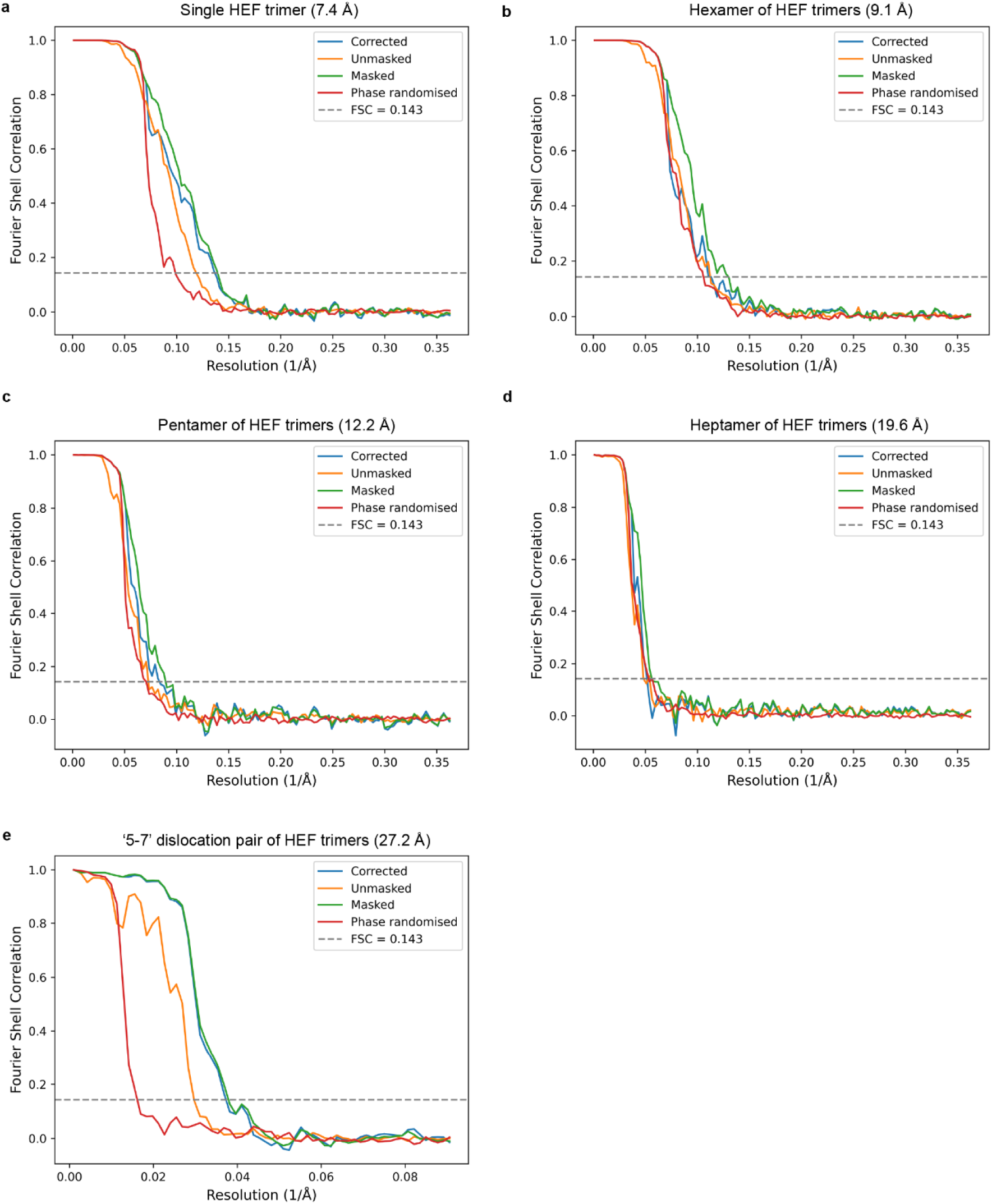
Fourier Shell Correlation (FSC) plot for all reconstructions. The curves are provided for the unmasked, masked, phase randomised, and corrected maps performed in RELION-3.0. The resolution is estimated at the FSC=0.143 threshold for the corrected curve. **a**, Single HEF trimer structure with a resolution of 7.4 Å. **b**, Hexamer of HEF trimers with a resolution determined at 9.1 Å. **c**, Pentamer of HEF trimers with a resolution determined at 12.2 Å. **d**, Heptamer of HEF trimers with a resolution determined at 19.6 Å. **e**, Composite oligomer comprising a “5-7” dislocation pair and an adjacent hexamer of HEF trimers with a resolution determined at 27.2 Å.

**Extended Data Table 1.** Data collection and processing parameters.

| Tomographic data collection | Dataset 1 | Dataset 2 | Dataset 3 |
| --- | --- | --- | --- |
| Virus growth method | MDCK cells | MDCK cells | Embryonated hen's eggs |
| Scheme | Bi-directional | Bi-directional | Bi-directional |
| Microscope | Titan Krios | Titan Krios | Titan Krios |
| Voltage (keV) | 300 | 300 | 300 |
| Tilt range (°) | -54 to +54 | -42 to +54/-54 to +54 | -42 to +54/-60 to +60 |
| Tilt interval (°) | 3 | 3 | 3 |
| Number of images per tilt series | 37 | 33/37 | 33/41 |
| Electron dose per tilt (e<sup>-</sup>/Å<sup>2</sup>) | 1.36 | 1.57 | 2.10 |
| Total electron dose per tilt series (e<sup>-</sup>/Å<sup>2</sup>) | 50.32 | 51.81/58.09 | 76/95 |
| Exposure rate (e<sup>-</sup>/px/s) | 0.85 | 7.6 | 4 |
| Detector | Falcon 3 | K2 Summit post GIF | K2 Summit post GIF |
| Calibrated pixel size (Å) | 2.23 | 2.20 | 1.38 |
| Defocus | -3.0 to -3.5 | -2.0 to -4.5 | -2.0 to -4.5 |
| Automation software | Tomography (FEI) | Tomography (FEI) | Tomography (FEI) |
| STA reconstruction |  |  |  |
| Number of tomograms used | 11 | 22 | 83 |
| Total number of extracted single HEF | 75828 | 120673 | 1100633 |
| Final subtomogram number of single HEF | 44275 | 50694 | 86446 |
| Final subtomogram number of pentameric HEFs | 914 | 2532 | 3108 |
| Final subtomogram number of hexameric HEFs | 29447 | 39182 | 35001 |
| Final subtomogram number of heptameric HEFs | 501 | 916 | 306 |
| Final subtomogram number of composite oligomer |  |  | 588 |
| Down-sampling rate | 2 | 4 | 1 |
| Resolution (Å, masked, FSC=0.143) |  |  |  |
| Single HEF | 12.7 | 17.7 | 7.4 |
| Pentameric HEF | 17.0 | 20.9 | 12.2 |
| Hexameric HEF | 13.7 | 20.9 | 9.1 |
| Heptameric HEF | 19.8 | 26.9 | 19.6 |
| Composite oligomer (bin 4) |  |  | 27.2 |

**Extended Data Table 2.**
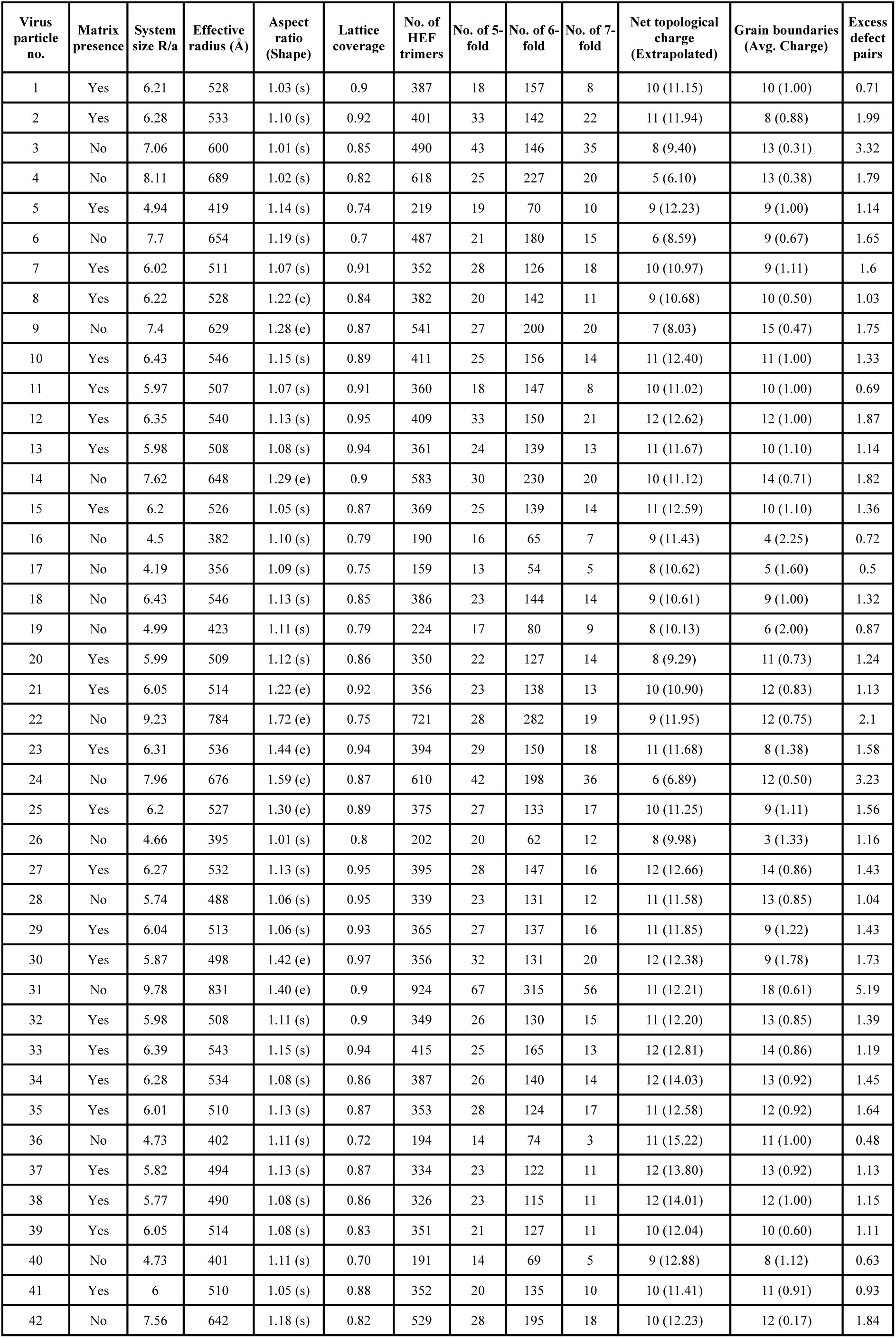
Parameters of the curated dataset of surface lattice models. A total of 42 virus particles of various shapes and sizes were modelled, with and without a matrix layer. The system size (*R/a*) is calculated by dividing area-equivalent radius *R* by lattice spacing *a* (∼85 Å) between adjacent HEF trimers. Virus shape, indicated in parentheses, was classified by aspect ratio: ellipsoidal (e) > 1.2 and spherical (s) ≤ 1.2. Lattice coverage was assessed by dividing the surface area of the incomplete surface lattice mesh to a fully closed mesh, with a mean coverage of 86 ± 7%. The total number of single HEF trimers and the centres of oligomers (pentamers, hexamers, and heptamers) are provided. These values were used to calculate the total topological charge and excess defect pairs per 12 disclinations presented in Fig. 4a**,c**, extrapolated for full surface coverage. The extrapolated mean total topological charge is 11.83 ± 1.52. Net topological charges were determined from observed defects, with extrapolated values shown in parentheses. The mean number of grain boundaries is 10.61 (12.33 ± 3.36 after extrapolation to full coverage), alongside the average net charge per grain boundary in parentheses with a mean of 0.96. Excess defect pairs scale with system size, as shown in Fig. 4c. Representative virus particles shown in Fig. 1 and Fig. 3 are particle no. 24, 16, 29, and 42.

## Data availability

Cryo-ET maps determined are deposited in the Electron Microscopy Data Bank (http://www.ebi.ac.uk/pdbe/emdb/, under accession numbers EMD-17001, EMD-17611, EMD-17612, and EMD-54338).

## Supplementary Information

**Supplementary Movie 1| Influenza C virus surface model showing the HEF glycoprotein lattice based on electron cryotomography.** The movie shows comparison of HEF oligomer maps determined by subtomogram averaging and their fitted coordinate models, informed by morphing from heptamer to hexamer to pentamer. Next, these oligomers are shown in a composite 5-6-7 arrangement in the context of an HEF surface lattice on a virus particle. The rotation of the surface model shows volume density of HEF trimers (grey) and spherical markers for hexamer (blue), pentamer (red), and heptamer (yellow) HEF oligomers and an underlying mesh describing the lattice. Next, the denoised tomogram is shown by scrolling through sections. The marker models and mesh are compared to the tomogram at the near and far surfaces of the virus particle.

## Acknowledgements

We thank A. Nans of the Structural Biology Science Technology Platform for assisting data collection; P. Walker and A. Purkiss of the Structural Biology Science Technology Platform; the Scientific Computing Science Technology Platform for computational support; and J. Molloy and J. Rubinstein for discussion. This work was supported by the Francis Crick Institute, which receives its core funding from the UK Medical Research Council (CC2106), Cancer Research UK (CC2106), and the Wellcome Trust (CC2106).

## Author contributions

Z.B.L. carried out research and analysed data. Z.B.L. and T.C. developed code to analyse the data. S.H. and L.J.C. performed experiments. Z.B.L. and P.B.R. conceived and designed research and wrote the paper with input from all authors.

## References

1. Hewat, E. A., Cusack, S., Ruigrok, R. W. H. & Verwey, C. Low resolution structure of the influenza C glycoprotein determined by electron microscopy. J Mol Biol 175, 175–193 (1984).

2. Flewett, T. H. & Apostolov, K. A reticular structure in the wall of influenza C virus. J Gen Virology 1, 297–304 (1967).

3. Halldorsson, S., Sader, K., Turner, J., Calder, L. J. & Rosenthal, P. B. In situ structure and organization of the influenza C virus surface glycoprotein. Nat. Commun. 12, 1694 (2021).

4. Bowick, M. J., Nelson, D. R. & Travesset, A. Interacting topological defects on frozen topographies. Phys Rev B 62, 8738–8751 (2000).

5. Bausch, A. R. et al. Grain Boundary Scars and Spherical Crystallography. Science 299, 1716–1718 (2003).

6. Caspar, D. L. D. & Klug, A. Physical Principles in the Construction of Regular Viruses. Cold Spring Harb Sym 27, 1–24 (1962).

7. Rosenthal, P. B. et al. Structure of the haemagglutinin-esterase-fusion glycoprotein of influenza C virus. Nature 396, 92–96 (1998).

8. Herrler, G. & Klenk, H.-D. Structure and Function of the Hef Glycoprotein of Influenza C Virus. Adv Virus Res 40, 213–234 (1991).

9. Muraki, Y. et al. A Mutation on Influenza C Virus M1 Protein Affects Virion Morphology by Altering the Membrane Affinity of the Protein▿. J Virol 81, 8766–8773 (2007).

10. Muraki, Y. et al. Identification of an amino acid residue on influenza C virus M1 protein responsible for formation of the cord-like structures of the virus. J Gen Virol 85, 1885–1893 (2004).

11. Zhang, X. et al. Structural and functional analysis of the roles of Influenza C virus membrane proteins in assembly and budding. J Biol Chem 101727 (2022) doi:10.1016/j.jbc.2022.101727.

12. Zhao, G. et al. Mature HIV-1 capsid structure by cryo-electron microscopy and all-atom molecular dynamics. Nature 497, 643–646 (2013).

13. Mattei, S., Glass, B., Hagen, W. J. H., Kräusslich, H.-G. & Briggs, J. A. G. The structure and flexibility of conical HIV-1 capsids determined within intact virions. Science 354, 1434– 1437 (2016).

14. Ganser, B. K., Li, S., Klishko, V. Y., Finch, J. T. & Sundquist, W. I. Assembly and Analysis of Conical Models for the HIV-1 Core. Science 283, 80–83 (1999).

15. Calcraft, T. et al. Integrated cryoEM structure of a spumaretrovirus reveals cross-kingdom evolutionary relationships and the molecular basis for assembly and virus entry. Cell (2024) doi:10.1016/j.cell.2024.06.017.

16. Cockburn, J. J. B. et al. Membrane structure and interactions with protein and DNA in bacteriophage PRD1. Nature 432, 122–125 (2004).

17. Kroto, H. W., Heath, J. R., O’Brien, S. C., Curl, R. F. & Smalley, R. E. C60: Buckminsterfullerene. Nature 318, 162–163 (1985).

18. Stone, A. J. & Wales, D. J. Theoretical studies of icosahedral C60 and some related species. Chem Phys Lett 128, 501–503 (1986).

19. Hashimoto, A., Suenaga, K., Gloter, A., Urita, K. & Iijima, S. Direct evidence for atomic defects in graphene layers. Nature 430, 870–873 (2004).

20. Kamien, R. D. The geometry of soft materials: a primer. Rev Mod Phys 74, 953–971 (2002).

21. Funkhouser, C. M., Sknepnek, R. & Cruz, M. O. de la. Topological defects in the buckling of elastic membranes. Soft Matter 9, 60–68 (2012).

22. Bowick, M., Cacciuto, A., Nelson, D. R. & Travesset, A. Crystalline Order on a Sphere and the Generalized Thomson Problem. Phys Rev Lett 89, 185502 (2002).

23. Thomson, J. J. On the structure of the atom: an investigation of the stability and periods of oscillation of a number of corpuscles arranged at equal intervals around the circumference of a circle; with application of the results to the theory of atomic structure. Philosophical Mag Ser 6 **7**, 237–265 (1904).

24. Wales, D. J. & Ulker, S. Structure and dynamics of spherical crystals characterized for the Thomson problem. Phys Rev B 74, 212101 (2006).

25. Dinsmore, A. D. et al. Colloidosomes: Selectively Permeable Capsules Composed of Colloidal Particles. Science 298, 1006–1009 (2002).

26. Irvine, W. T. M., Vitelli, V. & Chaikin, P. M. Pleats in crystals on curved surfaces. Nature 468, 947–951 (2010).

27. Lidmar, J., Mirny, L. & Nelson, D. R. Virus shapes and buckling transitions in spherical shells. Phys Rev E 68, 051910 (2003).

28. Zandi, R., Dragnea, B., Travesset, A. & Podgornik, R. On virus growth and form. Phys Reports 847, 1–102 (2020).

29. Brojan, M., Terwagne, D., Lagrange, R. & Reis, P. M. Wrinkling crystallography on spherical surfaces. Proc National Acad Sci 112, 14–19 (2015).

30. Baumeister, W. Cryo-electron tomography: A long journey to the inner space of cells. Cell 185, 2649–2652 (2022).

31. Martín-Bravo, M., Llorente, J. M. G., Hernández-Rojas, J. & Wales, D. J. Minimal Design Principles for Icosahedral Virus Capsids. Acs Nano 15, 14873–14884 (2021).

32. Zandi, R. & Reguera, D. Mechanical properties of viral capsids. Phys Rev E 72, 021917 (2005).

33. Zandi, R., Reguera, D., Bruinsma, R. F., Gelbart, W. M. & Rudnick, J. Origin of icosahedral symmetry in viruses. P Natl Acad Sci Usa 101, 15556–15560 (2004).

34. Seung, H. S. & Nelson, D. R. Defects in flexible membranes with crystalline order. Phys Rev A 38, 1005–1018 (1988).

35. Li, S., Roy, P., Travesset, A. & Zandi, R. Why large icosahedral viruses need scaffolding proteins. Proc National Acad Sci 115, 201807706 (2018).

36. Bowick, M. J., Cacciuto, A., Nelson, D. R. & Travesset, A. Crystalline particle packings on a sphere with long-range power-law potentials. Phys Rev B 73, 024115 (2006).

37. Einert, T., Lipowsky, P., Schilling, J., Bowick, M. J. & Bausch, A. R. Grain Boundary Scars on Spherical Crystals. Langmuir 21, 12076–12079 (2005).

38. Meng, G., Paulose, J., Nelson, D. R. & Manoharan, V. N. Elastic Instability of a Crystal Growing on a Curved Surface. Science 343, 634–637 (2014).

39. Guerra, R. E., Kelleher, C. P., Hollingsworth, A. D. & Chaikin, P. M. Freezing on a sphere. Nature 554, 346–350 (2018).

40. Burke, C. J., Mbanga, B. L., Wei, Z., Spicer, P. T. & Atherton, T. J. The role of curvature anisotropy in the ordering of spheres on an ellipsoid. Soft Matter 11, 5872–5882 (2015).

41. Vitelli, V., Lucks, J. B. & Nelson, D. R. Crystallography on curved surfaces. Proc National Acad Sci 103, 12323–12328 (2006).

42. Negro, G., Carenza, L. N., Gonnella, G., Marenduzzo, D. & Orlandini, E. Topological phases and curvature-driven pattern formation in cholesteric shells. Soft Matter 19, 1987– 2000 (2023).

43. Calder, L. J., Wasilewski, S., Berriman, J. A. & Rosenthal, P. B. Structural organization of a filamentous influenza A virus. Proc National Acad Sci 107, 10685–10690 (2010).

44. Beller, D. A. & Nelson, D. R. Plastic deformation of tubular crystals by dislocation glide. *Phys*. Rev. E 94, 033004 (2016).

45. Tanjeem, N., Wilkin, W. H., Beller, D. A., Rycroft, C. H. & Manoharan, V. N. Geometrical Frustration and Defect Formation in Growth of Colloidal Nanoparticle Crystals on a Cylinder: Implications for Assembly of Chiral Nanomaterials. ACS Appl. Nano Mater. 4, 10682–10691 (2021).

46. Zakharov, A. & Beller, D. A. Shape multistability in flexible tubular crystals through interactions of mobile dislocations. Proc. Natl. Acad. Sci. 119, e2115423119 (2022).

47. Mashl, R. J. & Bruinsma, R. F. Spontaneous-Curvature Theory of Clathrin-Coated Membranes. Biophys J 74, 2862–2875 (1998).

48. Cheng, Y., Boll, W., Kirchhausen, T., Harrison, S. C. & Walz, T. Cryo-electron Tomography of Clathrin-coated Vesicles: Structural Implications for Coat Assembly. J. Mol. Biol. 365, 892–899 (2007).

49. Kügelgen, A. von, Alva, V. & Bharat, T. A. M. Complete atomic structure of a native archaeal cell surface. Cell Reports 37, 110052 (2021).

50. Deatherage, J. F., Taylor, K. A., Amos, L. A. & Moody, M. F. Three-dimensional arrangement of the cell wall protein of Sulfolobus acidocaldarius. J. Mol. Biol. 167, 823–848 (1983).

51. Ben-Sasson, A. J. et al. Design of biologically active binary protein 2D materials. Nature 589, 468–473 (2021).

52. Crick, F. H. C. & Watson, J. D. Structure of Small Viruses. Nature 177, 473–475 (1956).

53. Chmielewski, D., Schmid, M. F., Simmons, G., Jin, J. & Chiu, W. Chikungunya virus assembly and budding visualized in situ using cryogenic electron tomography. Nat Microbiol 1–10 (2022) doi:10.1038/s41564-022-01164-2.

54. Azadi, A. & Grason, G. M. Neutral versus charged defect patterns in curved crystals. *Phys*. Rev. E 94, 013003 (2016).

55. Formanowski, F., Wharton, S. A., Calder, L. J., Hofbauer, C. & Meier-Ewert, H. Fusion Characteristics of Influenza C Viruses. J Gen Virol 71, 1181–1188 (1990).

56. Formanowski, F. & Meier-Ewert, H. Isolation of the influenza C virus glycoprotein in a soluble form by bromelain digestion. Virus Res 10, 177–192 (1988).

57. Mastronarde, D. N. & Held, S. R. Automated tilt series alignment and tomographic reconstruction in IMOD. J Struct Biol 197, 102–113 (2017).

58. Rohou, A. & Grigorieff, N. CTFFIND4: Fast and accurate defocus estimation from electron micrographs. J Struct Biol 192, 216–221 (2015).

59. Turoňová, B., Schur, F. K. M., Wan, W. & Briggs, J. A. G. Efficient 3D-CTF correction for cryo-electron tomography using NovaCTF improves subtomogram averaging resolution to 3.4Å. J Struct Biol 199, 187–195 (2017).

60. Bepler, T., Kelley, K., Noble, A. J. & Berger, B. Topaz-Denoise: general deep denoising models for cryoEM and cryoET. Nat Commun 11, 5208 (2020).

61. Liu, Y.-T. et al. Isotropic reconstruction for electron tomography with deep learning. Nat Commun 13, 6482 (2022).

62. Castaño-Díez, D., Kudryashev, M. & Stahlberg, H. Dynamo Catalogue: Geometrical tools and data management for particle picking in subtomogram averaging of cryo-electron tomograms. J Struct Biol 197, 135–144 (2017).

63. Zivanov, J. et al. New tools for automated high-resolution cryo-EM structure determination in RELION-3. Elife 7, e42166 (2018).

64. Bharat, T. A. M. & Scheres, S. H. W. Resolving macromolecular structures from electron cryo-tomography data using subtomogram averaging in RELION. Nat Protoc 11, 2054–2065 (2016).

65. Pettersen, E. F. et al. UCSF ChimeraX: Structure visualization for researchers, educators, and developers. Protein Sci 30, 70–82 (2021).

66. Krissinel, E. & Henrick, K. Inference of Macromolecular Assemblies from Crystalline State. J. Mol. Biol. 372, 774–797 (2007).

67. Chaillet, M. L., Roet, S., Veltkamp, R. C. & Förster, F. pytom-match-pick: A tophat-transform constraint for automated classification in template matching. J. Struct. Biol.: X 11, 100125 (2025).

68. Carr, J. C. et al. Reconstruction and representation of 3D objects with radial basis functions. Proc. 28th Annu. Conf. Comput. Graph. Interact. Tech. 67–76 (2001) doi:10.1145/383259.383266.

69. Taubin, G. Curve and surface smoothing without shrinkage. Proc. IEEE Int. Conf. Comput. Vis. 852–857 (1995) doi:10.1109/iccv.1995.466848.

70. Ermel, U. H., Arghittu, S. M. & Frangakis, A. S. ArtiaX: An electron tomography toolbox for the interactive handling of sub-tomograms in UCSF ChimeraX. Protein Sci 31, e4472 (2022).

71. Pedregosa, F. et al. Scikit-learn: Machine Learning in Python. JMLR 12, 2825--2830 (2011).

72. Kabsch, W. A solution for the best rotation to relate two sets of vectors. *Acta Crystallogr. Sect. A: Cryst. Phys., Diffr.*, Theor. Gen. Crystallogr. 32, 922–923 (1976).

73. Brunger, A. T. X-PLOR: Version 3.1: A System for x-Ray Crystallography and NMR. (Yale University Press, 1992).

74. Hendrickson, W. A. & Ward, K. B. A packing function for delimiting the allowable locations of crystallized macromolecules. Acta Crystallogr. Sect. A 32, 778–780 (1976).

75. Heymann, J. B., Chagoyen, M. & Belnap, D. M. Common conventions for interchange and archiving of three-dimensional electron microscopy information in structural biology. J Struct Biol 151, 196–207 (2005).

76. Muntoni, A. & Cignoni, P. PyMeshLab. (2021) doi:10.5281/zenodo.4438750.

77. Bernardini, F., Mittleman, J., Rushmeier, H., Silva, C. & Taubin, G. The ball-pivoting algorithm for surface reconstruction. Ieee T Vis Comput Gr 5, 349–359 (1999).

78. Nealen, A., Igarashi, T., Sorkine, O. & Alexa, M. Laplacian mesh optimization. Proc. 4th Int. Conf. Comput. Graph. Interact. Tech. Australas. Southeast Asia 381–389 (2006) doi:10.1145/1174429.1174494.

79. Boyé, S., Guennebaud, G. & Schlick, C. Least Squares Subdivision Surfaces. Comput. Graph. Forum 29, 2021–2028 (2010).

80. Koenderink, J. J. & Doorn, A. J. van. Surface shape and curvature scales. Image Vision Comput 10, 557–564 (1992).

81. Virtanen, P. et al. SciPy 1.0: fundamental algorithms for scientific computing in Python. Nat. Methods 17, 261–272 (2020).

82. Waskom, M., seaborn: statistical data visualization. J. Open Source Softw. 6, 3021 (2021).

